# Structural biases in disordered proteins are prevalent in the cell

**DOI:** 10.1101/2021.11.24.469609

**Authors:** David Moses, Karina Guadalupe, Feng Yu, Eduardo Flores, Anthony Perez, Ralph McAnelly, Nora M. Shamoon, Estefania Cuevas-Zepeda, Andrea D. Merg, Erik W. Martin, Alex S. Holehouse, Shahar Sukenik

## Abstract

Intrinsically disordered proteins and protein regions (IDPs) are essential to cellular function in all proteomes. Unlike folded proteins, IDPs exist in an ensemble of rapidly interchanging conformations. IDP sequences encode interactions that create structural biases within the ensemble. Such structural biases determine the three-dimensional shape of IDP ensembles and can affect their activity. However, the plasticity and sensitivity of IDP ensembles means structural biases, often measured *in vitro*, may differ in the dynamic and heterogeneous intracellular environment. Here we reveal that structural biases found *in vitro* in well-studied IDPs persist inside human-derived cells. We further show that a subset of IDPs are able to sense changes in cellular physical-chemical composition and modulate their ensemble in response. We propose that IDP ensembles can evolve to sense and respond to intracellular physicochemical changes, or to resist them. This property can be leveraged for biological function, be the underlying cause of IDP-driven pathology, or be leveraged for the design of disorder-based biosensors and actuators.

## Introduction

Intrinsically disordered proteins and protein regions (IDPs) play key roles in many cellular pathways and are vital to cellular function across all kingdoms of life^1,2^. Compared to folded proteins, IDPs lack a stable tertiary structure, have fewer intramolecular interactions, and expose a greater area of their sequence to the surrounding solution^3^. As a result, an IDP exists in an ensemble of conformations that can change rapidly in response to the physical-chemical characteristics of its surroundings^4,5^.

Despite being highly dynamic, IDP ensembles often contain structural biases, or preferences for certain subsets of conformations within the ensemble. Such structural biases may arise from short- or longer-range interactions within the protein sequence (**Fig. 1A**)^6^. An extensive body of work has established the importance of IDP structural biases to their function^2,7–12^. For example, local biases that form transient *ɑ*-helical segments modulate binding affinity in PUMA^9^ and p53^10,13^ or the liquid-liquid phase separation properties of TDP-43^12^. Changes to long-range structural biases were found to influence IDP function in p53^14^, BMAL1^15^ and Myc^16^. Thus, uncovering the structural biases of IDP ensembles is a prerequisite for understanding IDP function.

**Figure 1.**
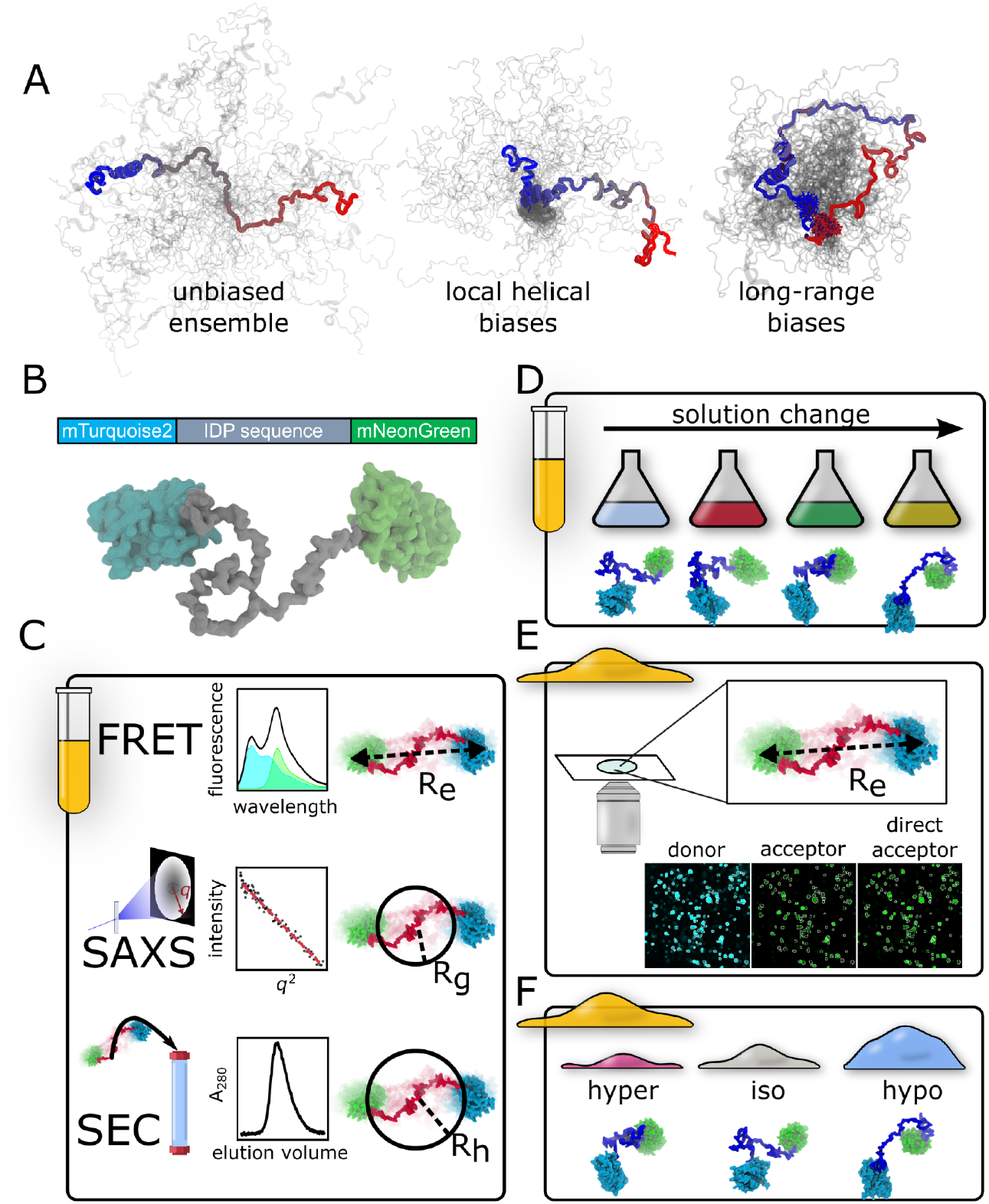
Schematic diagrams of methods used in this study. **(A)** IDP ensembles with and without structural biases. In all schemes, a single conformation is shown in color and other conformations are shown in gray. Structural biases increase the density in specific regions of the ensemble and alter its average dimensions. (**B**) FRET construct consisting of an IDP between two fluorescent proteins that serve as a FRET donor and a FRET acceptor. **(C)** *In vitro* experiments. Top: fluorescence resonance energy transfer (FRET). Middle: small angle X-ray scattering (SAXS). Bottom: analytic size-exclusion chromatography. **(D)** *In vitro* solution space scanning measures the FRET signal of a sequence in the presence of denaturing (urea, guanidinium), stabilizing (glycine, sarcosine), and crowding (PEG2k, Ficoll) solutes, as well as salt solutions (NaCl, KCl) that screen electrostatic interactions. **(E)** Live-cell FRET microscopy is performed on HEK293T cells expressing the same constructs used *in vitro*. **(F)** Changes in ensemble dimensions are measured in live cells following rapid hyperosmotic and hypoosmotic challenges.

Resolving IDP structural biases is especially important inside the heterogeneous cellular environment. Changes to the cellular physical-chemical composition occur regularly during the cell cycle^4,5,17,18^ (*e.g*., the breakdown of the nuclear envelope during mitosis^19^). Alternatively, these changes may result from pathology, such as the elevation in intracellular pH and rewiring of metabolic pathways common to nearly all cancer cells^20,21^. Such composition changes are known to affect even well-folded proteins^22–25^, but their effect on IDP structural biases has not been studied.

The structural malleability of IDP ensembles, coupled to the dynamic nature of the cellular environment, prompts two major unanswered questions: (1) To what degree are structural biases observed *in vitro* preserved inside the cell? (2) How do structural biases respond to the physical-chemical changes in the dynamic intracellular environment?

Here we aim to elucidate the structural biases of IDP ensembles in the cell, and understand how those biases change in response to physicochemical perturbations in the cellular environment. Our observations rely on ensemble fluorescence resonance energy transfer (FRET). To obtain a structural metric for IDP ensembles, we place sequences of interest between two FRET-pair fluorescent proteins (FPs), mTurquoise2 and mNeonGreen (**Fig. 1B**)^26–28^. Ensemble FRET provides, among other advantages, unmatched throughput and ease-of-use when working in live cells, but suffers from drawbacks when it comes to accurate quantification of distances. To mitigate these drawbacks, we have established a characterization pipeline that combines ensemble FRET (FRET, **Fig. 1C**), analytical size exclusion chromatography (SEC, **Fig. 1C**), small angle X-ray scattering (SAXS, **Fig. 1C**), changes in solution composition^4,28^ (**Fig. 1D**), and molecular simulations to identify structural biases of IDPs *in vitro*. We then leverage this characterization to examine the same constructs inside live cells using FRET microscopy (**Fig. 1E**). Finally, we perturb the cellular ensembles by subjecting cells to osmotic challenges that rapidly change cell volume, and measure the response of IDP ensembles through changes in FRET signal (**Fig. 1F**).

We first validate our pipeline using dipeptide Gly-Ser (GS) repeats, establishing these sequences as homopolymer benchmarks that contain no significant structural biases. We next compare the BH3 domain of PUMA, a naturally occurring IDP containing well-defined helical structural biases, against three variants where the wild-type sequence is scrambled. By scrambling a sequence, we confirm that structural biases are encoded in amino acid sequence, rather than amino acid composition. Finally, we investigate the structural ensembles of four well-studied IDPs inside live cells. We find that in all cases, the structural biases that define the ensemble *in vitro* also exist inside the cell. Furthermore, a subset of IDP sequences show a unique response to osmotically-triggered changes in cellular volume that was not observed in GS repeats.

Our work offers clear evidence that sequence-encoded structural biases exist in living cells, and further shows that these biases can be tuned by changes to the cellular environment. The existence of structural biases in IDP ensembles inside the cell suggests that they are subject to evolution, and can be rationally designed to create disorder-based sensors and actuators.

## Results

### Glycine-serine repeats are an unbiased, model-free standard to quantify IDP ensembles

Folded proteins are often compared to a standard (*e.g*., crystal) structure. For IDPs, no such standard exists. Instead, well-established homopolymer models are used as a reference^29,30^. No models exist for our dumbbell-shaped construct, especially not ones that are relevant in the cellular environment. We therefore wanted to create an empirical standard against which we can compare IDPs of arbitrary lengths.

As a benchmark against which to compare properties of naturally occurring heteropolymeric IDPs, we inserted homopolymeric dipeptide repeats into our FRET construct (**Fig. 2**). Specifically, we chose glycine-serine (GS) repeats for benchmarking because (1) they lack hydrophobicity, charge, and aromaticity which makes them easy to express and highly soluble^31^. (2) Previous studies have shown that GS-repeat sequences lack structural biases^32,33^ and (3) that they behave like ideal Gaussian chains in aqueous solutions^32,34,35^.

**Figure 2.**
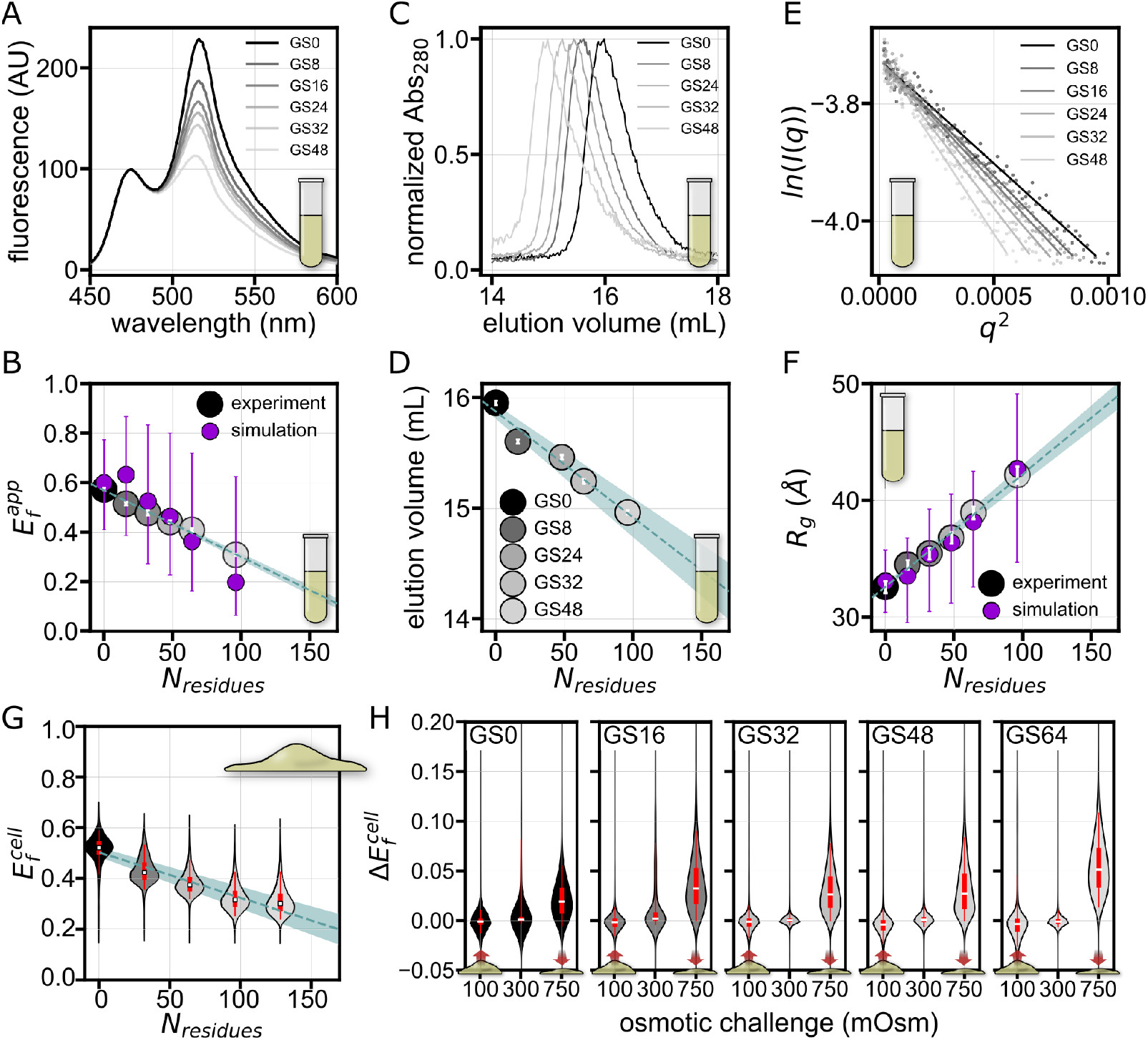
Characterization of GS repeat standards. **(A)** Fluorescence spectra from *in vitro* measurements of FRET GSX constructs, where X indicates the number of Gly-Ser repeats. **(B)** Apparent *in vitro* FRET efficiencies 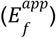 of GS repeats. Error bars represent error from two repeats. Here and in **Fig. 2D, 2F** and **2G**, dashed lines represent expected values for GS repeats of corresponding lengths based on a linear fit. Here and elsewhere, the blue shaded region represents the standard error of the linear fit. All-atom simulations of GS repeats are shown in purple, with error bars representing the median 50% of simulation results. **(C)** SEC chromatograms for GS repeats. **(D)** SEC elution volumes, expressed as the position of the peak in mL, vs. number of residues in the GS-repeat sequence. Error is assumed to be one frame in each direction. **(E)** Guinier regions and fitted lines from SAXS experiments for GS repeats. **(F)** Radii of gyration (*R*_*g*_), derived from Guinier analysis of SAXS data for GS repeats. White error bars represent error from fitting lines to Guinier plots. The same all-atom simulations of GS repeats shown in **Fig. 2B** are used to calculate the simulated *R*_*g*_, shown in purple, with purple error bars representing the median 50% of simulation results. *R*_*g*_ values from the pairwise distribution function (*P(r)*) for all IDPs in this study are shown in **Table S1. (G)** FRET efficiencies of GS-repeats measured in live cells 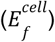 In all live-cell data, violin plots span the entire dataset and their thickness represents the probability. The median is shown as a white square, and the median 50% and 95% are shown as thick and thin lines at the center of the violin, respectively. The blue line is a linear fit of the medians, and fit errors shown by the shaded region. **(H)** Response to osmotic challenge expressed as change in 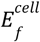 before and after the challenge 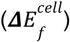. The dataset used to generate all of the live-cell figures is shown as **Table S2**. For the number of cells used to generate each violin plot see **Table S3**.

Ensemble FRET experiments provide an apparent FRET efficiency 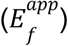, which is inversely proportional to the distance between the two FPs. When 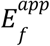 is high FPs are close together (compact), and when low they are far apart (expanded). As previously reported, 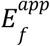 decreased linearly with the number of GS repeats in a dilute buffer solution^5^ (**Fig. 2A,B, S1**). However, the three-dimensional structure of the ensemble cannot be resolved by a single distance measurement^36–38^. To obtain additional, orthogonal measurements that can inform about the structure of the ensemble, we performed size-exclusion chromatography coupled with small-angle X-ray scattering (SEC-SAXS) on the same constructs used for FRET^39–41^. The chromatograms obtained from SEC showed a consistent, linear size-dependent increase in elution volume (**Fig. 2C,D, S2**), indicating that the proteins increase in dimension with GS repeat length. Analysis of SAXS intensity curves showed a similar linear dependence on GS length (**Fig. 2E,F, S3, S4**), displaying linearly increasing radii of gyration (*R*_*g*_, **Fig. 2F**) in agreement with our other results. Finally, we conducted all-atom simulations of all GS repeats. Our simulations assumed that the FPs only take up space (*i.e*., are non-interacting) and that GS repeats behave like homopolymers. From these simulations, ensembles were selected to quantitatively match the SAXS scattering data (**Fig. S5**). These ensembles reproduced the GS length-dependent 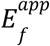 values as well, indicating the simulation conditions at least managed to reproduce our experimental results (**Fig. 2B,F**). Together, all methods consistently show the same length-dependent trend for the GS repeats, and that the length of the construct, rather than *e.g*. intramolecular interactions between or with FPs, is the dominant factor affecting these dimensions. The excellent quantitative agreement with our simulations further indicates that GS repeats behave like ideal homopolymers, which lack structural biases^36,37^.

To further verify that GS repeats do not contain structural biases, we conducted FRET-based solution space scanning of GS-repeat constructs^4,28^. Solution space scanning probes structural biases in the ensemble by modulating interactions between the sequence and the solution. We reason that if structural biases exist, different GS repeat lengths will show a different structural response to the same solution. We measure the change in FRET efficiency 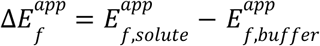 for all GS repeat lengths in a range of solution conditions (**Fig. S6**). Our scan showed that GS repeats of all lengths responded identically to polymeric crowders, denaturants and stabilizing osmolytes (**Fig. S6**). Overall, the internal consistency of the results from our orthogonal characterization methods establishes GS repeats as a model-free, homopolymer standard which lacks structural biases.

### Live-cell measurements recapitulate *in vitro* results for GS repeat ensembles

We next sought to establish GS repeats as a bias-free standard in live cells. To facilitate direct and straightforward comparison with our *in vitro* experiments, we used the same genetically encoded FRET constructs as we had used *in vitro*. GS-repeat FRET constructs were expressed in HEK293T cells, which all showed similar morphology and expression levels regardless of the construct being expressed (**Fig. S7**).

Our live-cell measurements of GS repeats showed trends in FRET efficiency calculated from live-cell imaging 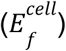 that are in quantitative agreement with *in vitro* measurements (**Fig. 2B,G**). Notably, in live cells our FRET constructs show a much wider variability in 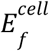 compared to 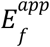 *in vitro*. This variability may be caused by a wide range of factors, including cell-to-cell differences in composition, cell state, and differences in construct expression levels. Despite this, the remarkable agreement with *in vitro* data indicates that the lack of structural biases for GS repeats detected *in vitro* persists inside live cells.

To test whether GS ensemble dimensions are sensitive to the cellular environment, we subjected cells to osmotic challenges. To resolve their immediate effects on a protein, such perturbations should be performed rapidly and measured as quickly as possible to prevent any kind of transcriptional response^42,43^. To this end, we use rapid osmotic challenges induced by the addition of NaCl (hyperosmotic, 750 mOsm) or water (hypoosmotic, 100 mOsm) to media (300 mOsm). Osmotic challenges were previously shown to produce robust and reproducible changes in cellular volume through the efflux or influx of water^42–44^. We report on the difference in FRET signal of each cell following this perturbation, 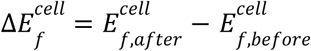. The measurements before and after the challenge are collected within a span of 45 seconds or less (**Fig. 1F**).

Hyperosmotic perturbations resulting in cell shrinkage caused a positive 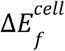 that scaled with the length of the construct (**Fig. S8**). This is in line with previous studies of IDPs in crowded conditions and in the cell^18,30,44^, and can be explained by the increased ability of longer sequences to compact. Hypoosmotic perturbations, on the other hand, produced no significant change in 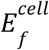 (**Fig. S8**). This lack of response was surprising, especially considering the fact that GS polymers are capable of expansion *in vitro* (**Fig. S6**). Regardless, our osmotic challenge experiments define a standard for the response of bias-free IDP ensembles to osmotically induced changes in cellular volume.

### Amino acid sequence determines IDP structural biases and their response to changes in solution composition

Having established a reliable homopolymer standard *in vitro* and in live cells, we set out to investigate how a naturally occurring IDP compares with our GS repeats. We chose the wildtype sequence of the PUMA BH3 domain (WT or PUMA WT) (**Fig. 3A,B**) because its functionally important residual helicity is a well-studied example of local structural bias in IDPs^9,45^. We first established the previously reported short-range helical structural biases of the unlabeled sequence^46^ as indicated by the characteristic double minima in the circular dichroism (CD) spectrum (**Fig. 3B,C**). Next, we measured the 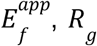, and SEC elution volume of WT PUMA using our *in vitro* pipeline (WT in **Fig. 3D-F**). Although in SEC WT PUMA eluted near the same volume as would be expected of GS repeats of the same length (**Fig. 3E**), SAXS and FRET showed WT PUMA to be significantly more compact than corresponding GS repeats (**Fig. 3D,F**), confirming that we are able to detect local structural biases present in WT PUMA but absent in GS repeats.

**Figure 3.**
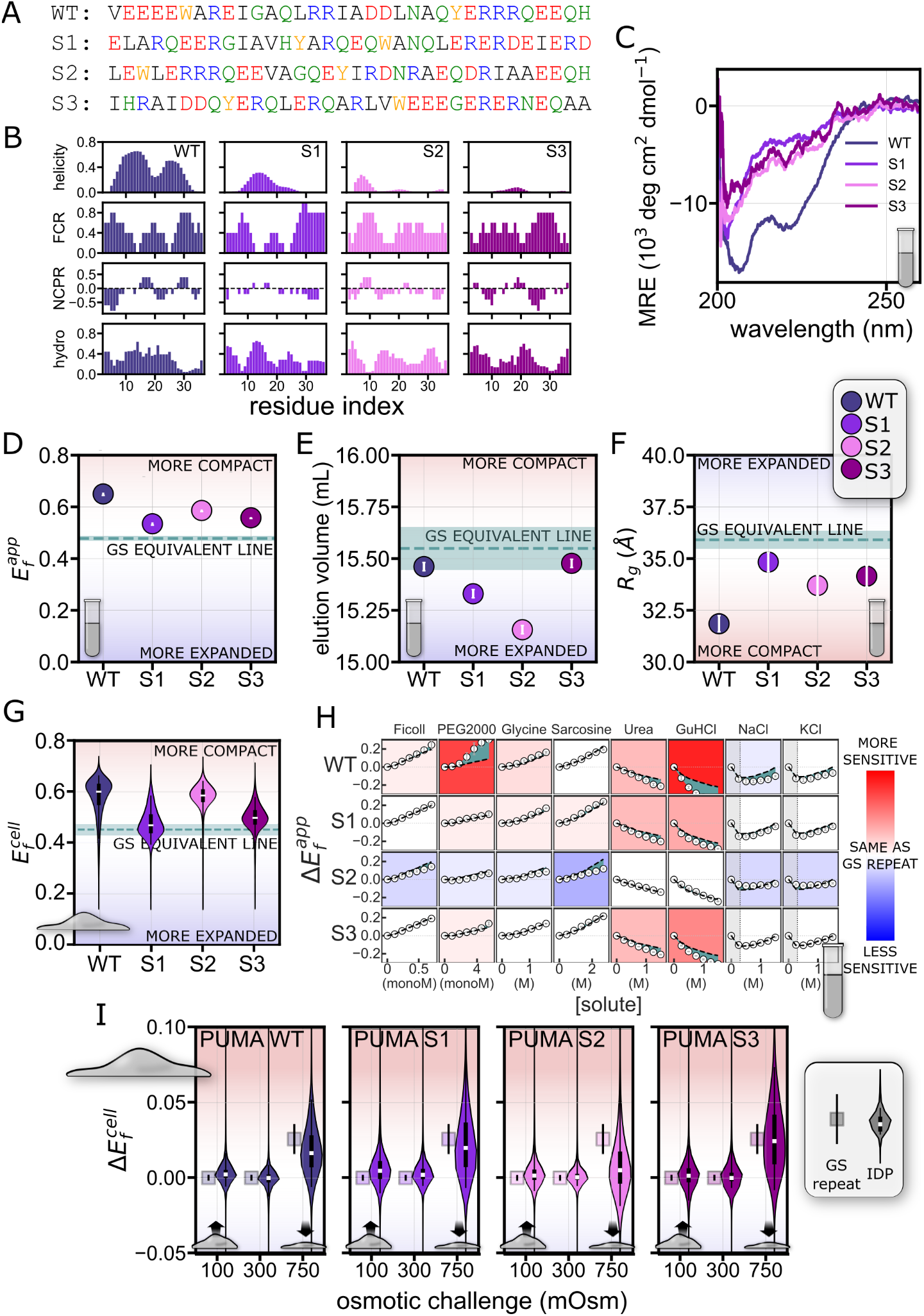
**(A)** Sequence of wild-type PUMA BH3 domain (WT PUMA) and three sequences (S1, S2, S3) derived by shuffling WT PUMA’s sequence. Red: negative charge; blue: positive charge; black: hydrophobic residues; green: polar residues; orange: aromatic residues. **(B)** Molecular features of WT PUMA and shuffles. Predicted helicity was calculated using all-atom simulations. Other parameters were evaluated by localCIDER^49^. FCR: fraction of charged residues. NCPR: net charge per residue. hydro: Kyte-Doolittle hydrophobicity. Value at each position on the x axis represents the average of the indicated residue and its four nearest neighbors. **(C)** Circular dichroism (CD) spectroscopy signatures of PUMA variants without flanking FPs, taken at concentrations same as FRET experiments. CD experiments at other concentrations are shown in **Fig. S9. (D)** 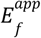 of PUMA constructs. Error bars represent error from two repeats. Here and in **Fig. 3E-G**, dashed line represents the expected value for a GS-repeat construct of the same length (34 residues) as WT PUMA and scrambles (GS-equivalent). **(E)** SEC elution volume at peaks for PUMA constructs. Error is assumed to be one frame in each direction. **(F)** *R*_*g*_ of PUMA constructs. Error bars represent error from fitting lines to Guinier plots. **(G)** 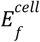 of PUMA constructs. Features are as in **Fig. 2G. (H)** Solution space scans of PUMA constructs, with results expressed as 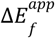, the difference between 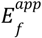 of an IDP construct in a given solution condition and in a dilute buffer. White dots: 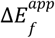 of IDP. Black dashed lines: interpolated 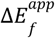 of a GS repeat of the same length as the IDP (**Fig. S12**). Blue-green shaded regions between white dots and black dashed lines: difference between 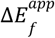 of IDP and GS repeat. Heatmap backgrounds: red shows more sensitivity (more expansion or compaction) than a GS repeat of the same length; blue shows less sensitivity than a GS repeat; white shows the same sensitivity as a GS repeat; deeper shades show greater difference in sensitivity from GS repeats. Shaded regions on left side of cells for solutes NaCl and KCl: approximate range of concentrations within which electrostatic screening is the dominant effect; the leftmost two points of each series, since they are within that range, are not used in the assignment of background color. **(I)** Osmotic challenge of HEK293T cells expressing PUMA constructs. Violin plots represent the data for PUMA constructs and squares represent 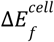 GS-repeat equivalent. Features are as in **Fig. 2G,H**. For the number of cells used to generate each violin plot see **Table S3**.

Is residual helicity like that observed in WT PUMA a prerequisite for detectable structural biases? To answer this question, we generated sequence scrambles of WT PUMA (**Fig. 3A, Table S1**) and measured their ensembles *in vitro*. Sequence scrambles retain the amino acid composition but disrupt any structural biases that may be present^37,38^. We generated three scrambles of WT PUMA with different patterning of charged and hydrophobic residues (S1, S2, and S3, **Fig. 3A,B**). To test the existence of helical structural biases in the scrambled sequences, we measured the secondary structure of the label-free IDPs using CD. As expected, the CD spectra of the scrambles showed no double minima (**Fig. 3C, S9**), indicating that the helical structural biases of WT PUMA were no longer present.

We next characterized ensemble dimensions for all scrambles using FRET (**Fig. 3D**), SEC (**Fig. 3E**), SAXS (**Fig. 3F**), and all-atom Monte Carlo simulations (**Fig. S10**). FRET and SAXS show that not only are the scrambles more compact than a GS repeat of the same length, they also all differ from each other, despite having similar CD spectra and identical amino acid composition (**Fig. 3A-C**). The overall agreement between trends from FRET and SAXS measurements shows that the WT PUMA sequence ensemble is significantly the most compact, followed by more closely-grouped S2, S3, and finally S1. This trend is also recapitulated in label-free, all-atom simulations, indicating that tethering to the two fluorescent protein labels does not change the trends in ensemble dimension for this measurement (**Fig. S10**). SEC data shows a different trend, with all sequences appearing more expanded than a GS linker and S3 showing an almost equal compaction to the WT (**Fig. 3E**). This may be due to chemical interactions between the constructs and the SEC column matrix.^47^ However, since all sequences contain the same amino acid composition, even these different interactions indicate structuring within the ensemble. The differences shown in all methods, not only between WT PUMA and the three scrambles but also between each scramble, highlight that structural biases exist even in the absence of the helical structural biases in the WT sequence. Our measurements highlight that the WT PUMA ensemble is uniquely more compact than the scrambles.

We hypothesized that different structural biases in PUMA and its scrambles would also manifest in their response to different solutions. To test this, we performed solution space scans for all four PUMA variants (**Fig. S11**). We report these results as 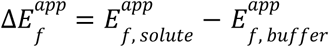, and compare these to the interpolated 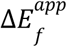 of GS repeats of the same length in the same solution condition (**Fig. 3H, S12**). Deviations from 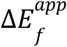 of length-equivalent GS repeats indicate higher/lower sensitivities of the sequences (indicated by red/blue backgrounds, respectively) (**Fig. 3H**). We were surprised to find that despite having the most compact ensemble, WT PUMA showed the highest sensitivity of all scrambles (as indicated by the stronger blue and red backgrounds in **Fig. 3H**). Specifically, the WT sequence displayed stronger compaction in response to polymeric crowders (specifically PEG2000) and stronger expansion in response to denaturants (urea and GuHCl) compared to both the corresponding GS-repeat sequence and the three sequence scrambles. The three scrambles showed milder responses, with a notable difference for S2, which was significantly less sensitive to all solutes (as indicated by the blue background). These differences indicate that IDPs possess sensitivity to the chemical composition of their environment that is encoded in their sequence. Furthermore, the presence of structural biases does not preclude ensemble sensitivity to the surrounding solution, and may even amplify it.

### Sequence-dependent structural biases seen *in vitro* persist in live cells, but *in vitro* solution responses are not necessarily recapitulated

We next wanted to see if the structural biases measured *in vitro* for WT PUMA and its scrambles were retained in the cellular environment. We expected helical structural biases to persist in the cell due to the relatively extensive interaction network that forms them^48^, but reasoned that biases within scrambled sequences were weaker and therefore might not be retained. To test this, we performed our live-cell FRET imaging experiments on WT PUMA and the three variants (**Fig. 3G**). We found striking agreement with the FRET measurements done in dilute aqueous buffers (**Fig. 3D**). Specifically, both the relative magnitude and the trend in *E*_*f*_ measured *in vitro* was replicated in live cells, with WT > S2 > S3 > S1. Overall, 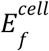 reveals that the structural biases found in these sequences *in vitro* persist inside the cell, even in the absence of short-range helical structural biases (which occur only in WT PUMA).

Our next goal was to measure whether these ensembles differ in their response to changes in the cellular environment. We again used osmotically-triggered cell volume perturbations as a means to reproducibly change the concentration of all cellular solutes. The change in value of each cell’s average FRET signal before and after the osmotic challenge, 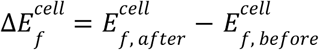 is reported and compared with the expected 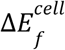 for a GS-repeat equivalent (**Fig. 3I**). We were surprised to find that the WT sequence, which displayed more sensitivity than a corresponding GS-repeat sequence to certain solutes *in vitro*, showed a response similar to that of GS repeats under both cell volume increase and decrease. Remarkably, this similarity to GS-repeat sensitivity in live cells was seen in all sequences except S2, which displayed a significantly lower tendency than the other three sequences to compact under hyperosmotic conditions. The lower sensitivity of S2 was also observed *in vitro* (**Fig. 3H**). This result indicates that IDP ensemble sensitivity to changes in the cellular environment is encoded in sequence, but is difficult to predict since it may or may not correlate with the sensitivity measured in dilute buffers.

### Structural biases are prevalent in naturally occurring IDPs

Having seen that structural biases seen *in vitro* persist inside the cell for PUMA and its scrambles, we wanted to see whether this is a general property of other IDP sequences. We inserted a range of well-studied IDPs of different lengths into our construct and characterized them *in vitro* and in live cells. We tested the N-terminal activation domain (NTAD) of p53 (residues 1-61, p53)^10^, the low-complexity domain of FUS (residues 1-163, FUS)^50^, the N-terminal region of the adenovirus hub protein E1A (residues 1-40, E1A)^51^, and the C-terminal region of the yeast transcription factor Ash1 (residues 418-500, Ash1)^52^ (**Fig. S13, Table S1**). Here, comparison to a GS-repeat equivalent becomes even more critical to facilitate a length-independent comparison between constructs.

Using our *in vitro* characterization pipeline, we found clear divergence in nearly all constructs from GS repeats. Our FRET experiments show that all but two sequences are much more compact than a GS-repeat sequence of the same dimensions (indicated by 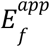 values above the GS line, **Fig. 4A**). The two that fell close to the GS line, p53 and Ash1, were indeed reported to be relatively expanded in other studies^10,52^. A similar trend was observed for SAXS-derived *R*_*g*_ values (**Fig. 4C**). SEC data (**Fig. 4B**) shows mostly similar trends, though PUMA, E1A, and p53 appear to be more expanded than GS repeats. As before, the deviations from the GS-equivalent line, together with the changes in trends between characterization methods, highlight the differences in structural biases between different IDP sequences.

**Figure 4.**
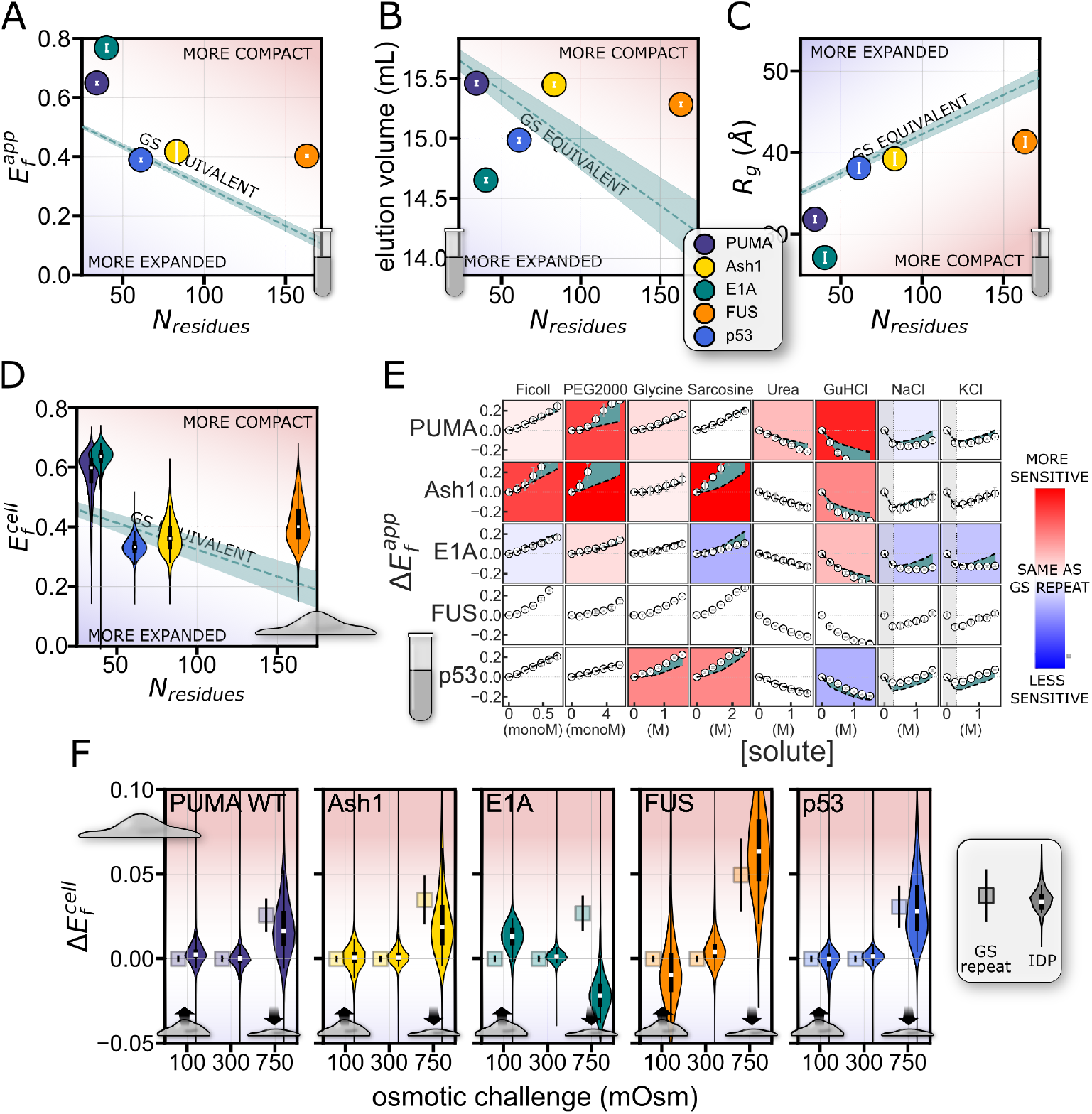
Comparison of global dimensions and solution sensitivity of GS repeats and naturally occurring IDPs. **(A)** 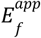 of IDR constructs. Error is from two repeats. Here and in **Fig. 4B-D**, dashed line represents expected values for GS repeat sequences of corresponding lengths. **(B)** SEC elution volume of IDR constructs. Error is assumed to be one frame in each direction. **(C)** *R*_*g*_ of IDR constructs. Errors are from fitting lines to Guinier plots. **(D)** 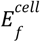 of IDR constructs. Features are as in **Fig. 2G. (E)** Solution space scans of IDR constructs. Features are as in **Fig. 3H**, except for FUS for which the GS repeat trend could not be accurately extrapolated. **(F)** Osmotic challenge of IDR constructs. Features are as in **Fig. 2G,H**. For the number of cells used to generate each violin plot, see **Table S3**.

Our next goal was to determine the extent to which the structural biases observed *in vitro* for these constructs persist in the cell. As before, we expressed the same constructs in HEK293T cells, and used live-cell imaging to quantify the in-cell FRET efficiency, 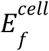 (**Fig. 4D**). This time, results are compared to GS repeats of the equivalent length, indicated by a dashed line. Remarkable agreement was observed between 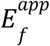 measured *in vitro* and the in-cell 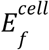 values (**Fig. 4A,D**). One exception is E1A, which showed a more compact conformation *in vitro*, but became more expanded inside the cell. This agreement indicates that the structural biases for these IDPs that determined their ensemble shape *in vitro* largely exist inside the cellular environment. E1A is an example where this rule does not hold. Furthermore, compaction inside the cell cannot be explained by macromolecular crowding^25,30,53^, and indicates that some sequences may have other factors (*e.g*., protein-protein interactions) that define ensemble shape.

### Naturally occurring IDPs differ in their sensitivity to solution changes

To ascertain whether similar structural metrics to GS repeats in a dilute buffer solution imply a complete lack of structural preferences, we performed solution space scanning on each of these IDP constructs^5^ (**Fig. 4E, S14**). As expected, different sequences showed markedly different sensitivities to the solutes used. PUMA and Ash1 showed an outlying degree of sensitivity, with significantly larger changes compared to GS repeats of the same length in both compacting and expanding solutes (**Fig. S6, S12**), while E1A appeared to be less sensitive to the same solutes. The response to salts also showed deviations, with significantly less response to high salt concentrations for E1A (indicated by the constant value in 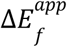 at concentrations above 0.25 mM salt). Interestingly, p53, whose dimensions were closest to those of its GS equivalent in dilute buffer (**Fig. 4A**), also displayed the closest sensitivity to its GS equivalent (**Fig. 4E**). This wide range of responses to changes in solution conditions agrees with our expectations based on our FRET, SAXS, and SEC results, further supporting the existence of sequence-dependent structural biases in the ensembles of naturally occurring IDPs. Moreover, the different IDP ensembles show differing and specific sensitivities to changes in their chemical environment.

Finally, we wanted to measure the response of these IDPs to changes in intracellular composition. We subjected cells to hypoosmotic or hyperosmotic challenges and followed the changes in average FRET signal for each cell, 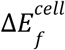 (**Fig. 4F**). We compare these to the changes expected for GS repeats of the same length, shown as the squares adjacent to each violin plot. We observe that some sequences behaved as expected from GS repeats—namely PUMA, Ash1, and p53 all fall within the range expected of a GS-repeat equivalent. FUS displayed a similar behavior to GS repeats upon hyperosmotic challenge, but showed an outlying ability to expand in hypoosmotic conditions. However, the most striking results were shown by E1A, which compacted when the cell expanded, and expanded when the cell compacted. This counterintuitive trend stands in stark contrast to most *in vitro* observations of monomeric IDPs (with some notable exceptions^54^), and cannot be explained by macromolecular crowding of the cellular environment^25^. We therefore propose that the behavior of E1A could be the result of protein-protein interactions or post-translational modifications that occur inside the cell.

Taken together, these results show not only that structural biases in IDP ensembles exist both *in vitro* and inside the cell, but also that IDP ensembles detect and respond to changes in the composition of their environment. This ability to respond to physical-chemical changes is encoded in sequence, and occurs both in a test tube and in the cell. However, despite the agreement between IDP structural biases in a dilute solution *in vitro* and in isosmotic conditions in the cell, comparing *in vitro vs* in-cell solution sensitivity is not straightforward: We see general agreement between *in vitro* and in-cell behavior, especially in terms of compaction when the solution is crowded (using PEG2000 or Ficoll) or the cell is compacted, but also intriguing exceptions. One such exception is PUMA, whose ability to resist change is weak *in vitro* but strengthened in the cellular environment. And E1A’s mild sensitivity to composition changes *in vitro* makes its counterintuitive behavior of expanding when the cell compacts all the more striking. These distinctive behaviors suggest a link to function. How does PUMA’s ability to resist change in the cellular environment contribute to its role in apoptosis? How does the expansion of the adenovirus E1A IDP in a compacting cell relate to its ability to bind host proteins? Future studies into these questions may help reveal the mechanistic underpinnings of function and malfunction in these and other IDPs in the cellular environment.

### Limitations and drawbacks

One drawback of this work is the use of fluorescent proteins (FPs) in our constructs. There are many advantages to genetically encoded FRET constructs. They can be produced easily in *E. coli* with no need for further labeling, or transiently or stably expressed in any genetically tractable cell line. Additionally, they assist with sequence solubility, increase signal from scattering methods, and hinder phase separation in the case of FUS. However, the presence of bulky, folded domains tethered to the IDP of interest may affect our results through the intramolecular interaction of the FPs with themselves or with the IDP sequence. We cannot, and do not, claim that the FPs in these constructs are inert. Nonetheless, concerns regarding artifacts from our use of FPs are mitigated by (1) the use of the same FPs for all constructs and the comparison against GS-repeat constructs, which facilitate meaningful comparison between all sequences; (2) the agreement between our experiments and all-atom simulations of IDPs which do not include FP interactions (**Fig. 1B,F, S10**); (3) if different IDPs tethered to the same FPs cause different IDP-FP or FP-FP interactions, it is still an indication of differing IDP structural biases which persist inside the cell; and (4) the fact that our IDPs of interest (and those in many other IDP studies) are tethered to folded domains in their native state. Overall, the self-consistency of our dataset, and the careful comparison to GS repeats, indicate that intramolecular FP-FP or FP-IDR interactions are not the main driver of the differences observed here, and do not negate our main conclusions.

## Discussion

The study of disordered proteins requires shifting from the classical sequence-structure-function paradigm to one where the structural biases of the ensemble beget function^31^. While an extensive body of work has established the existence of structural biases in IDP ensembles *in vitro*, few studies have attempted to do so in the cell. Our results systematically show that structural biases *in vitro* are prevalent in IDP sequences, are encoded in amino acid sequence rather than composition, and exist even in the absence of local secondary structural biases (*e.g*., local helical preference, **Fig. 1A**). The cell is often treated as a chemically monolithic environment, yet spatial and temporal regulation of volume, water content, pH, ions, and metabolites accompany key processes and pathology in cell biology^55–57^. Our in-cell study establishes that IDP structural biases observed *in vitro* occur in live cells for almost all cases reported here. Furthermore, both in cells and *in vitro*, IDP structural biases can be rewired in response to changes in the cellular environment. This provides a mechanistic explanation for numerous cases where IDPs sense and actuate a response to such changes^58–60^: a change in structural bias in response to physical-chemical changes can alter IDP function. Importantly, sensing and actuating through this mechanism occurs at the speed of protein conformational changes and requires no additional energy (*e.g*., ATP).

The encoding of IDP ensemble sensitivity in amino acid sequence suggests that it is subjected to evolutionary selection. We propose that certain sequences have evolved to act as sensors and actuators of changes in the cellular environment. In the same vein, other disordered sequences have the ability to resist structural changes (as shown for the case of PUMA S2). Indeed, changes in ensemble structure provide a rapid, specific, and energetically efficient way for IDPs to sense and respond to changes in the cellular environment. As our understanding of IDP sensing expands, we expect to uncover novel functions for this important class of proteins. In addition, learning to predict and control this sensitivity will allow for the design of IDP-based sensors targeting specific physicochemical intracellular conditions, as has already been demonstrated for the case of osmotic pressure sensing^60^.

An additional implication of the evolved ability to sense and respond to changes in the environment is that a misregulated intracellular environment may disparately affect IDP function. Metabolic rewiring, a hallmark of cancer, viral infection, and other pathologies, can dramatically alter the physicochemical composition of the cell^61,62^. Even if this change would alter the activity of only a small subset of IDPs, their role as central signaling hubs could cause widespread cellular malfunction. In this way, IDP sequences can be drivers of pathology in a deleterious cellular environment, even in the absence of mutations. We propose that this phenomenon is a previously overlooked cause of IDP-driven proteopathies.

## Supplementary data

Supplementary **Figures S1-S20** and **Table S4** are available in Supplementary Information. Supplementary **Tables S1-S3** are available as csv files. Source data and code to produce all figures in this manuscript are available online at https://github.com/sukeniklab/IDP_structural_bias.

## Acknowledgements

We thank J.A. Caro, M. Gebala, A. LiWang, J. Riback, P.S. Romero-Pérez, H.B. Schmidt, and M. Thompson for helpful comments and discussion. We are indebted to J. Hopkins, S. Chakravarthy, and all BioCAT beamline staff at the Advanced Photon Source at Argonne National Laboratory, I. Rajkovic and all BioSAXS beamline staff at SLAC National Accelerator Laboratory, and G. Hura and all staff at the SIBYLS beamline at Lawrence Berkeley Laboratory for assistance with SAXS measurements. We thank M. Sadqi of the Center for Cellular and Biomolecular Machines (CCBM) for help with mass spectrometry. Research reported in this publication was supported by the NIH under award R35GM137926 to SS. KG is supported by a fellowship from NSF-CREST Center for CCBM at UC Merced, Grant No. NSF-HRD-1547848. This research used the Advanced Photon Source at Argonne National Laboratory under Contract No. DE-AC02-06CH11357, Proposal No. 75514. We acknowledge computing time on the MERCED cluster at UC Merced, NSF Grant ACI-1429783, and on the XSEDE computational infrastructure framework, Grant No. TG-MCB190103 to ASH and SS, supported by NSF Grant ACI-154856.

## Materials and Methods

### FRET construct design and cloning

The FRET backbone for bacterial expression (fIDP_pET-28a(+)-TEV) or for mammalian expression (fIDP_pCDNA3.1(+)) was prepared by ligating mTurquoise2 and mNeonGreen into pET28a-TEV or pCDNA backbone using 5’ NdeI and 3’ XhoI restriction sites. Genes encoding for IDP regions were obtained from GenScript (Piscataway, NJ) and ligated between the two fluorescent proteins using 5’ SacI and 3’ HindIII restriction sites. Cloned plasmids were amplified in XL1 Blue (Invitrogen) cell lines using manufacturer-supplied protocol. Sequences of all IDP inserts are available in **Table S1**.

### FRET construct expression and purification

BL21 (DE3) cells were transformed with fIDP_pET-28a(+)-TEV plasmids according to manufacturer protocol and grown in LB medium with 50 μg/mL kanamycin. Cultures were incubated at 37 °C while shaking at 225 rpm until OD600 of 0.6 was reached (approx. 3 h), then induced with 1 mM IPTG and incubated for 20 h at 16 °C while shaking at 225 rpm. Cells were harvested by centrifugation for 15 min at 3,000 rcf, the supernatant was discarded, and the cells were lysed in lysis buffer (50 mM NaH_2_PO_4_, pH 8, 0.5 M NaCl) using a QSonica Q700 Sonicator (QSonica, Newtown, CT). Lysate was centrifuged for 1 h at 20,000 rcf and the supernatant collected and flowed through a column packed with Ni-NTA beads (Qiagen). The FRET construct was eluted with 50 mM NaH_2_PO_4_, pH 8, 0.5 M NaCl, 250 mM imidazole, and further purified using size-exclusion chromatography on a Superdex 200 PG column (GE Healthcare) in an AKTA go protein purification system (GE Healthcare). The purified FRET constructs were divided into 200 μL aliquots, flash-frozen in liquid nitrogen, and stored at -80 °C in 20 mM sodium phosphate buffer, pH 7.4, with the addition of 100 mM NaCl. Protein concentration was measured after thawing and before use using UV-vis absorbance at 434 and 506 nm (the peak absorbance wavelengths for mTurquoise2 and mNeonGreen, respectively; the molar absorbance coefficients for mTurquoise2 and mNeonGreen are 30,000 cm^-1^M^-1^ and 116,000 cm^-1^M^-1^, respectively^63^. Calculations of concentration based on λ = 434 nm produced slightly higher values than calculations based on λ = 506 nm, so the concentrations based on the measurement at λ = 506 nm were used), and purity was assessed by SDS-PAGE after thawing and before use. To verify the brightness of the FPs, we measured the UV-Vis absorbance of both donor and acceptor molecules before each FRET assay. We used only samples that displayed an absorbance ratio Abs_506_/Abs_434_ = ratio of 2.8 ± 0.2, a reasonable ratio given the difference in the molar extinction coefficients of mTurquoise2 and mNeonGreen. Samples where the ratio deviated from this value were discarded.

### Preparation of solutions for solution-space scanning

Solutes were purchased from Alfa Aesar (Sarcosine, PEG2000), GE Healthcare (Ficoll), Thermo Scientific (Guanidine Hydrochloride), and Fisher BioReagents (Glycine, Potassium Chloride, Sodium Chloride, Urea), and used without further purification. Stock solutions were made by mixing the solute with 20 mM sodium phosphate buffer, pH 7.4, with the addition of 100 mM NaCl except for experiments where the concentration of NaCl and KCl were varied, which began free of additional salt. The same buffer was used for all dilutions.

### *In vitro* FRET experiments

*In vitro* FRET experiments were conducted in black plastic 96-well plates (Nunc) with clear bottom using a CLARIOstar plate reader (BMG LABTECH). Buffer, stock solution, and purified protein solution were mixed in each well to reach a volume of 150 μL containing the desired concentrations of the solute and the FRET construct, with a final concentration of 1 μM protein. Fluorescence measurements were taken from above, at a focal height of 5.7 mm, with gain fixed at 1020 for all samples. For each FRET construct, two repeats from different expressions with 6 or 12 replicates each were performed in neat buffer, and two repeats from different expressions were done in every other solution condition. Fluorescence spectra were obtained for each FRET construct in each solution condition by exciting the sample in a 16 nm band centered at λ = 420 nm, with a dichroic at λ = 436.5 nm, and measuring fluorescence emission from λ = 450 to 600 nm, averaging over a 10 nm window moved at intervals of 0.5 nm. Base donor and acceptor spectra for each solution condition were obtained using the same excitation and emission parameters on solutions containing 1 μM mTurquoise2 or mNeonGreen alone, and measuring fluorescence emission from 450 to 600 nm^63,64^.

### Calculation of FRET efficiencies and end-to-end distances

The apparent FRET efficiency 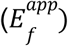 of each FRET construct in each solution condition was calculated by linear regression of the fluorescence spectrum of the FRET construct with the spectra of the separate donor and acceptor emission spectra in the same solution conditions (in order to correct for solute-dependent effects on fluorophore emission). 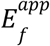 was calculated using the equation^65^:

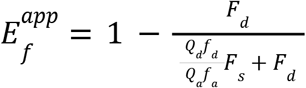

where *F*_*d*_ is the decoupled donor contribution, *F*_*s*_ is the decoupled acceptor contribution, *f*_*d*_ is the area-normalized donor spectrum, *f*_*a*_ is the area-normalized acceptor spectrum, *Q*_*d*_ = 0.93 is the quantum yield of mTurquoise2, and *Q* = 0.8 is the quantum yield of mNeonGreen^26,64^.

The data for each series of solution conditions consisting of increasing concentrations of a single solute was processed in the following manner:

1. Raw spectra for the free donor and free acceptor in the various solution conditions were loaded, and the averages of all repeats in each solution condition were computed. These averages are referred to as the “raw” donor and acceptor spectra below because they will be further corrected.
2. The donor and acceptor peak intensities were assumed to change in a linear fashion with increasing solute concentration, so peak height of donor- or acceptor-only spectra vs. concentrations were linearly fit.
3. To correct for artifacts (such as variations in FRET construct concentration between different wells) that may contribute to unexpected differences in fluorescence intensity, a correction factor was applied to each raw donor and acceptor spectrum to bring the peak intensity to the linear fit described in step 2, resulting in “corrected” donor and acceptor spectra. Importantly, we have seen in our previous work that this correction corrects well-to-well variations in raw data but has a negligible effect on overall values and trends^5^.
4. The raw FRET construct fluorescence spectra for the series were loaded.
5. To compensate for unintended direct excitation of the acceptor by excitation at the donor excitation frequency, the corrected acceptor spectrum for each solution condition was subtracted from the FRET construct spectrum for each solution condition, resulting in “corrected” FRET construct spectra.
6. The corrected donor, acceptor and FRET construct spectrum for each solution condition was fitted with a linear regression function to determine the decoupled contributions of the donor and acceptor to the FRET construct spectrum.
7. 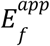 of each FRET construct in each solution condition was calculated using the equation shown above.

### Size exclusion chromatography and small-angle X-ray scattering experiments

Small-angle X-ray scattering (SAXS) experiments were performed at BioCAT (beamline 18ID at the Advanced Photon Source, Chicago). The experiments were performed with in-line size exclusion chromatography (SEC-SAXS) (**Fig. S2**) to separate monomeric protein from aggregates and improve the accuracy of buffer subtraction. Experiments were conducted at 20 °C in 20 mM sodium phosphate, pH 7.4, with 100 mM NaCl. Samples of approximately 300 µL were loaded, at concentrations in mg/mL approximately equal to 240 divided by the molecular weights of the constructs in kD (for example, a typical construct of molecular weight 60 kD would have a target concentration for SEC-SAXS of 240/60 = 4 mg/mL), onto a Superdex 200 Increase 10/300 column (GE Life Sciences) and run at 0.6 mL/min using an ÄKTA Pure FPLC system (Cytiva). The column eluent passed through the UV monitor and proceeded through the SAXS flow cell which consists of a 1.5 mm ID quartz capillary with 10 μm walls. The column to X-ray beam dead volume was approximately 0.1 mL. Scattering intensity was recorded using a Pilatus3 1M (Dectris) detector placed 3.5 m from the sample providing access to a q-range from 0.003-0.35 Å^-1^. 0.5 second exposures were acquired every 2 seconds during the elution. Data was reduced at the beamline using BioXTAS RAW version 2.1.1^66,67^. The contribution of the buffer to the X-ray scattering curve was determined by averaging frames from the SEC eluent which contained baseline levels of integrated X-ray scattering, UV absorbance and conductance. Frames were selected as close to the protein elution as possible and, ideally, frames pre- and post-elution were averaged. Multiple peaks for GS48, WT PUMA, E1A, and FUS were deconvolved using evolving factor analysis (EFA) (**Fig. S15**)^68,69^ and the peak with calculated molecular weight corresponding to the monomer was chosen for further analysis. Final scattering profiles were generated by subtracting the average buffer trace from all elution frames and averaging curves from elution volumes close to the maximum integrated scattering intensity; these frames were statistically similar in both small and large angles. Buffer subtraction and subsequent Guinier fits (**Fig. S3**), as well as Kratky transformations (**Fig. S4**), deconvolution of peaks using EFA, molecular weight calculations based on volume of correlation^70^ and Porod volume^71^ (**Table S1**), and pair distance distribution (P(r)) analysis using the indirect Fourier transform (using the algorithm in the GNOM program by Svergun and Semyenuk) were done in BioXTAS RAW. Radii of gyration (*R*_*g*_) were calculated from the slope of the fitted line of the Guinier plot at maximum *q* x *R*_*g*_ = 1 using the equation^72^:

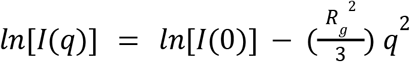

### Mammalian cell culture

HEK293T cells were cultured in Corning treated flasks with Dulbecco’s modified Eagle medium (Advanced DMEM:F12 1X, Gibco Cat. No. 12634-010) supplemented with 10% FBS (Gibco Cat. No. 16000-044) and 1% penicillin/streptomycin (Gibco Cat. No. 15140-122). For live-cell microscopy experiments, 5,000 cells were plated in a µ-Plate 96-well black treated imaging plate (Ibidi Cat. No. 89626) and allowed to adhere overnight (∼16 hours) before transfection. Cells were incubated at 37 °C and 5% CO_2_. Before transfection, the media was switched out with new warmed media. XtremeGene HP (Sigma Cat. No. 6366236001) was used to transfect FRET construct plasmids into HEK293T cells per manufacturer’s protocol. Cells were incubated at 37 °C and 5% CO_2_ for 48 hours. NaCl stock solution was prepared by dissolving NaCl (Fisher Bioreagents CAS 7647-14-5) in 1X PBS (Gibco Cat. No. 70011-044) and filtering using a 0.2 µm filter. The solutions used for perturbations were obtained by diluting the imaging media (1X PBS) with autoclaved DI water to achieve hypoosmotic (100 mOsm) conditions or by adding NaCl stock solution for hyperosmotic (750 mOsm) conditions. Isoosmotic (300 mOsm) conditions were obtained by adding 1X PBS. To prepare for imaging, cells were rinsed once with 1X PBS and left in 200 μL PBS (300 mOsm) for imaging.

### Live-cell microscopy

Imaging was done on a Zeiss epifluorescent microscope using a 10X 0.3 NA dry objective. Excitation was done with a Colibri LED excitation module and data was collected on a duocam setup with two linked Hamamatsu Flash v3 sCMOS cameras. The cells were imaged at room temperature before and after perturbation with 150 ms exposure times. Imaging was done by exciting mTurquoise2 at 430 nm (donor and acceptor channels, **Fig. 1E**) or mNeonGreen at 511 nm (direct acceptor channel, **Fig. 1E**). Emitted light was passed on to the camera using a triple bandpass dichroic (467/24, 555/25, 687/145). When measuring FRET, emitted light was split into two channels using a downstream beamsplitter with a 520 nm cutoff. For each perturbation, the cells were focused using the acceptor channel and imaged before manually adding water (hypoosmotic condition), PBS (isosmotic condition) or NaCl solution (hyperosmotic condition) and pipetting up and down 10 times to ensure mixing. The final osmolarities that were used for the perturbations were: 100 mOsm, 300 mOsm (isosmotic), and 750 mOsm with NaCl as the osmotic agent. Imaging was typically completed in ∼ 45 seconds.

### Image analysis

Images were analyzed using ImageJ^73^. Images collected before and after osmotic challenge, containing three channels each, were stacked and aligned using the StackReg plugin with rigid transformation (**Fig. S16**).^74^ The aligned image was segmented based on the donor channel before perturbation. Segmentation was done using several methods to ensure that the results were robust. The methods included the ImageJ built-in implementations of the Triangle and MinError algorithm, as well as a fixed threshold that selected only pixels with intensities between 1,500 - 40,000. All methods gave nearly identical results, so the fixed threshold method was finally selected for the data shown in all live cell figures. The resulting mask was processed using the Open and Watershed binary algorithms of imageJ. Cells were selected using the Analyze Particles option of ImageJ, picking only those with an area between 65 - 845 μm^2^, and with a circularity of 0.1 - 1.0. The resulting regions of interest were averaged in each channel at each timepoint. The resulting cells were filtered to remove cells with an intensity over 10,000 (to correlate with *in vitro* experiment concentrations, see **Fig. S17**) and cells where the absolute change in direct acceptor emission was over 2,000 (which tended to be cells that moved or lifted off the coverslip during measurement). To correct for donor bleedthrough and cross-excitation, cells were transfected with the mTurquoise2 or mNeonGreen construct only, the cells were imaged and analyzed using the same protocol as previously mentioned, and correlation plots were generated to determine percent bleedthrough and cross-excitation (**Fig. S18**). The final filtering step removed cells with a corrected donor/acceptor ratio that was negative or higher than 6. Cell FRET efficiency before and after perturbation (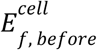 and 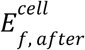 respectively) was calculated by 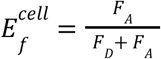. The resulting dataset is available as **Table S2**. The number of cells measured for each construct and condition from this dataset are summarized in **Table S3**. Analysis code is available as an ImageJ macro at https://github.com/sukeniklab/IDP_structural_bias.

### *In vitro* concentration dependence experiments

Protein aliquot samples were diluted into a series of varying concentrations using 20 mM sodium phosphate, 100 mM NaCl, pH 7.4 buffer. Samples were prepared on a µ-Plate 96-well black treated imaging plate (Ibidi Cat. No. 89626). Fluorescent beads (Phosphorex Cat. No. 2225) were added to the prepared aliquots to ensure focus on the bottom of the well. Imaging parameters were the same parameters as were used for the live-cell microscopy experiments. Images were also analyzed using ImageJ^73^. Instead of segmentation, the center of the images were selected and the average pixel intensities were measured. In order to correlate emission with concentration, we plotted protein concentration against direct acceptor emission (**Fig. S17**).

### Label-free peptide synthesis and purification

WT PUMA and shuffled sequences were prepared via standard microwave-assisted solid-phase peptide synthesis protocols using a Liberty Blue automated microwave peptide synthesizer (CEM, NC, USA) and ProTide Rink Amide resin (CEM). Fmoc-deprotection was achieved by treatment with 4-methylpiperidine (20% v/v) in dimethylformamide (Sigma-Aldrich), and Fmoc-amino acids were activated using N,N’-Diisopropylcarbodiimide (Sigma-Aldrich) and Oxyma Pure (CEM). Peptides were N-terminally acetylated and C-terminally amidated. After synthesis, the peptidyl resins were filtered and rinsed with acetone and air-dried. The crude peptides were cleaved from the resin for 4 hours at room temperature with a 92.5% trifluoroacetic acid (TFA), 2.5% H_2_O, 2.5% 3,6-dioxa1,8-octane-dithiol, 2.5% triisopropylsilane cleavage solution, precipitated with cold diethyl ether, and centrifuged at 4000 rpm for 10 min at 4 °C. After centrifugation, the supernatants were discarded, and the pellets were dried under vacuum overnight. Crude peptides were purified by high-performance liquid chromatography (HPLC) using an Agilent 1260 Infinity II HPLC instrument equipped with a preparative scale Phenomenex Kinetex XB-C18 column (250 × 30 mm, 5 μm, 100 Å) (**Fig. S19**). Peptides were eluted with a linear gradient of acetonitrile-water with 0.1% TFA. The target fractions were collected, rotovapped, and lyophilized. Purified peptides were analyzed by mass spectrometry using a Q-Exactive Hybrid Quadrupole-Orbitrap mass spectrometer (Thermo Scientific) (**Fig. S20, Table S4**).

### CD spectroscopy

Lyophilized protein constructs were weighed and dissolved in a 20 mM sodium phosphate, 100 mM NaCl buffer at pH 7.4 to make a 200 μM stock. The stock was diluted into a concentration series to measure the CD spectra. CD spectra were measured using a JASCO J-1500 CD spectrometer with a 1 cm quartz cell for 1 μM and 2 μM protein concentration and 0.1 cm quartz cell for other concentrations (Starna Cells, Inc., Atascadero, CA) using a 0.1 nm step size, a bandwidth of 1 nm, and a scan speed of 200 nm/min between 260 to 190 nm. Each spectrum was measured 7 times and averaged to increase the signal-to-noise ratio. The buffer control spectrum was subtracted from each protein spectrum. CD spectra were normalized using UV 280 nm absorbance to eliminate the small concentration difference between different protein constructs.

### All-atom simulations of constructs with fluorescent proteins

To verify that our *in vitro* SAXS and FRET results report on the same conformational ensemble, we performed all-atom simulations of full-length constructs that include both fluorescent proteins using an identical amino acid sequence to the experimental constructs. Fluorescent protein models were constructed from PDB files 4AR7 (mTurquoise2)^75^ and 5LTR (mNeonGreen)^76^. Simulations were performed using the ABSINTH implicit solvent model and CAMPARI Monte Carlo simulation engine^77^.

Considering the size of these proteins, simulating them at full-length and all-atom resolution raises a number of challenges. Given that our objective here was to determine whether SAXS and FRET were in agreement in the context of a simple homopolymeric linker, we took advantage of the ABSINTH implicit forcefield’s ability to tune specific components of the Hamiltonian. Specifically, we performed simulations in which all excluded-volume interactions were present (*i.e*., the repulsive component of the Lennard-Jones potential was turned on). However, the attractive component of the Lennard-Jones potential was only turned on for residues within the glycine-serine (GS) linker, and limited only to intra-linker interactions by varying the inherent Lennard-Jones parameters of all atoms outside of the GS linker. Beyond these two components, all additional non-bonded Hamiltonian terms (*i.e*., long and short-range electrostatics and solvation effects) were turned off, dramatically lowering the computational cost of simulations. By systematically tuning the overall strength of the attractive GS repeat intramolecular interactions, we in effect performed simulations for GS homopolymers for all relevant homopolymer interaction strengths and GS-repeat lengths from 0 to 48 (*i.e*., 0 residues to 96 residues).

We initially performed simulations using a GS0 construct, where the only backbone degrees of freedom available were associated with the set of flexible residues that connect the two beta-barrels. Specifically, all backbone dihedral angles for amino acids within the two beta-barrels were switched off, but all sidechain degrees of freedom were accessible. The residues between the two beta-barrels that had their backbone degrees of freedom sampled consist of amino acids 227 to 255 (GITLGMDELYKEGLSKLMVSKGEEDNMAS) in the GS0 construct^5^. After running thousands of short independent simulations in which these twenty-nine amino acids were sampled with variable intramolecular interaction strengths, we subselected an ensemble of 1000 distinct conformations which, on average, reproduced the experimentally measured SAXS scattering data for the GS0 construct (**Fig. S5A**). This GS0 ensemble was then used to define the starting configurations of the FPs and the ‘handles’ (non-GS component of the construct) for all other GS simulations.

For each of the other GS-repeat lengths (8, 16, 24, 32, 48), we performed simulations in which the attractive Lennard-Jones potential was scaled from 0.30 to 0.62 in steps of 0.02. This range straddles intramolecular interaction strengths that cause the longer GS chains to behave as a self-avoiding random coil (attractive LJ scaling parameter = 0.3) and a compact globule (attractive LJ scaling parameter = 0.62). For each combination of GS length and LJ strength, we performed 1000 independent simulations in which the fluorescent proteins and associated handles defined in the GS0 simulations were also fixed in place. As such, in total we performed 17,000 independent simulations for each separate GS length (i.e., 85,000 independent simulations in total). This approach enables to in effect construct a large collection of ensembles with (defined GS lengths but variable GS intramolecular interaction strengths) from which we will ultimately subselect and concatenate many individual simulation trajectories based on the agreement between the simulated scattering profile and the real scattering profile. These data will be used to construct a sub-ensemble that recapitulates the experimental scattering data - i.e. an unbiased, data-driven approach to construct an ensemble consistent with the experimental measurements.

Each simulation was run in a spherical droplet with a radius of 500 Å to avoid any possible finite size effects. Given the absence of any electrostatic components, no ions were included in the simulations. Each simulation was run for 100,000 Monte Carlo steps. The first 50,000 steps were discarded as equilibration, and conformations were then sampled every 5000 steps. As such, each independent simulation generated 10 conformations, such that each GS/LJ combination generated a 10,000 conformer ensemble (1000 independent simulations with 10 conformations per simulation). Other than the repulsive component of the Lennard-Jones potential and (for some atoms) the attractive component of the Lennard-Jones potential, all other modes of nonbonded interactions were switched off. As such, each individual simulation takes on the order of 10 minutes.

Having performed this set of simulations we calculated predicted scattering profiles for each independent simulation using FoXS software, as described previously^78,79^. To assess the agreement between each short simulation and the experimental scattering data we computed 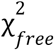, a parameter explicitly developed to assess the goodness-of-fit for scattering data^70^].We calculated 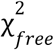 for GS-length matching simulations to assess how well each length-matched sub-ensemble compared to the experimentally measured scattering data. Using a 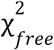 of 3.2 (a large value that reflects the relatively small error in the experimentally measured SAXS data), we generated sub-ensembles with scattering curves that quantitatively reproduced the experimental data at each of the GS-repeat lengths (**Fig. S5A**).

Finally, using the SAXS-matched sub-ensembles, we computed the inter-barrel distance based on the distance between two residues in the center of the beta-barrel (**Fig. S5B**). Distances were calculated between alpha-carbon atoms, such that we subtracted a 6 Å offset to approximately account for the distance between the alpha-carbon atoms and the anticipated chromophore centers. The resulting inter-beta-barrel distances are in excellent agreement with distances measured from ensemble FRET experiments. For **Fig. 2B**, these end-to-end distances (*R*_e_) were converted to *E*_f_ values using the equation 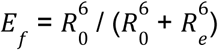 assuming *R*_0_ the Förster distance for the mTurquoise2-mNeonGreen FRET pair, to be 62 Å^64^. Taken together, this approach shows that the ensembles that best describe the SAXS data also correctly describe the distances inferred from FRET, confirming that these orthogonal methods are reporting on the same underlying conformational ensemble. The final sub-ensembles for each GS-repeat length and the associated data are provided in https://github.com/sukeniklab/IDP_structural_bias. Simulation analysis was performed with SOURSOP (https://soursop.readthedocs.io/).

### All-atom simulation of IDP region only

Simulations of label-free IDP sequences used in this study were done using the CAMPARI simulation suite and the ABSINTH forcefield^77,80^. For each sequence, five independent simulations were run at 310 K using 8×10^7^ Monte Carlo steps (following 1×10^7^ steps of equilibration) starting from random conformations to ensure proper sampling. Protein conformations were written out every 12,500 steps. The end-to-end distance and the helicity of the simulated conformation ensembles were determined using the MDTraj python library^81^.

## Notes

https://github.com/sukeniklab/IDP_structural_bias

### Competing Interest Statement

The authors have declared no competing interest.

### Summary of Updates

Now showing in-cell data for 4 additional IDPs. The narrative has been changed to focus on the similarity of IDP ensembles in vitro and in cells. All new text and figures.

