## Supplemental Information for "Structural biases in disordered proteins are prevalent in the cell"

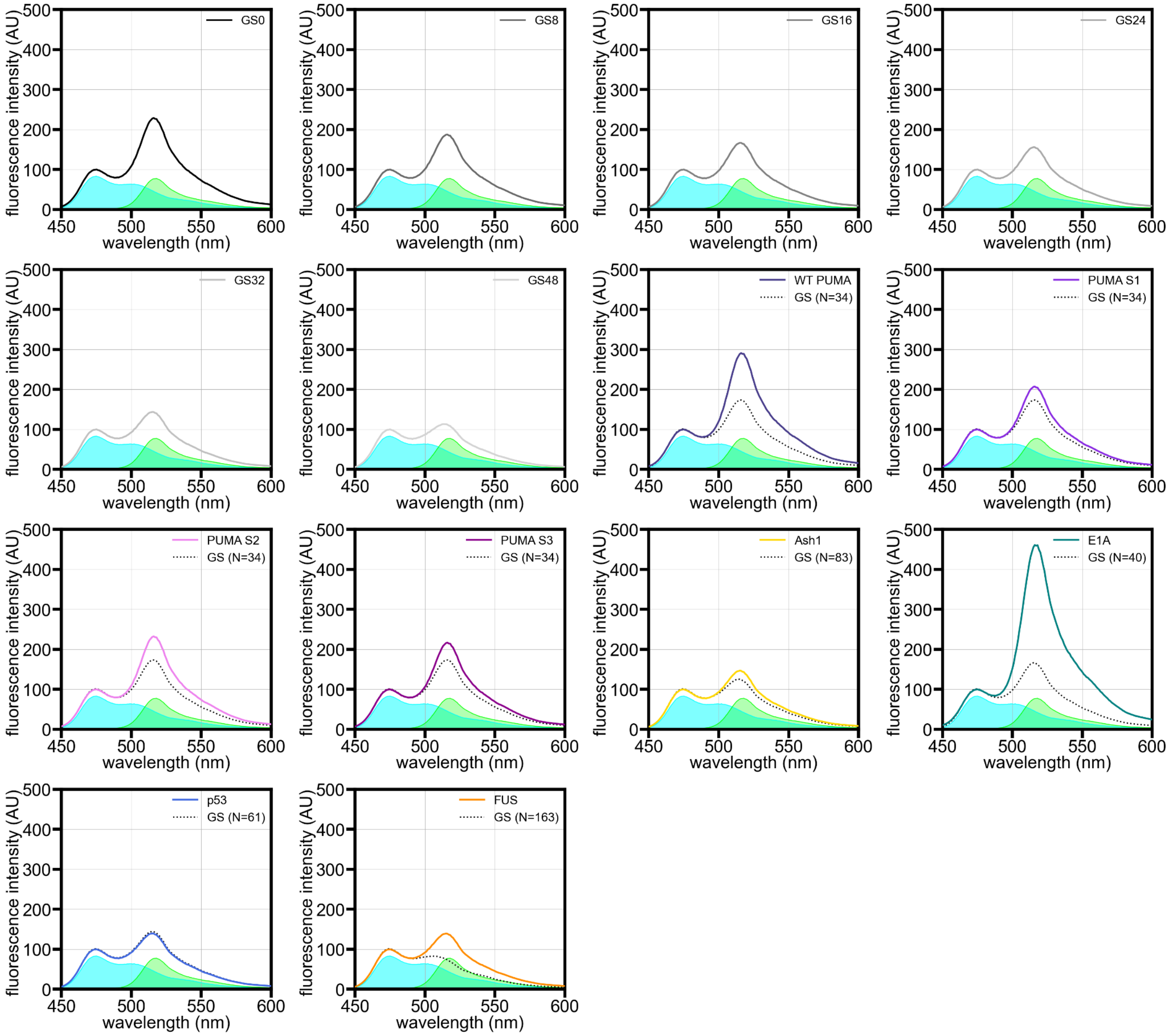


**Figure S1.** Fluorescence spectra from measurements of FRET constructs in a dilute phosphate buffer solution. Spectra of constructs incorporating IDRs that are not GS-repeat sequences are compared with expected fluorescence spectra of FRET constructs incorporating GS-repeat sequences of equal length (black dotted curves), where N refers to the number of amino acids. Blue and green shaded areas are the base spectra for mTurquoise2 and mNeonGreen, respectively, in the same buffer solution.

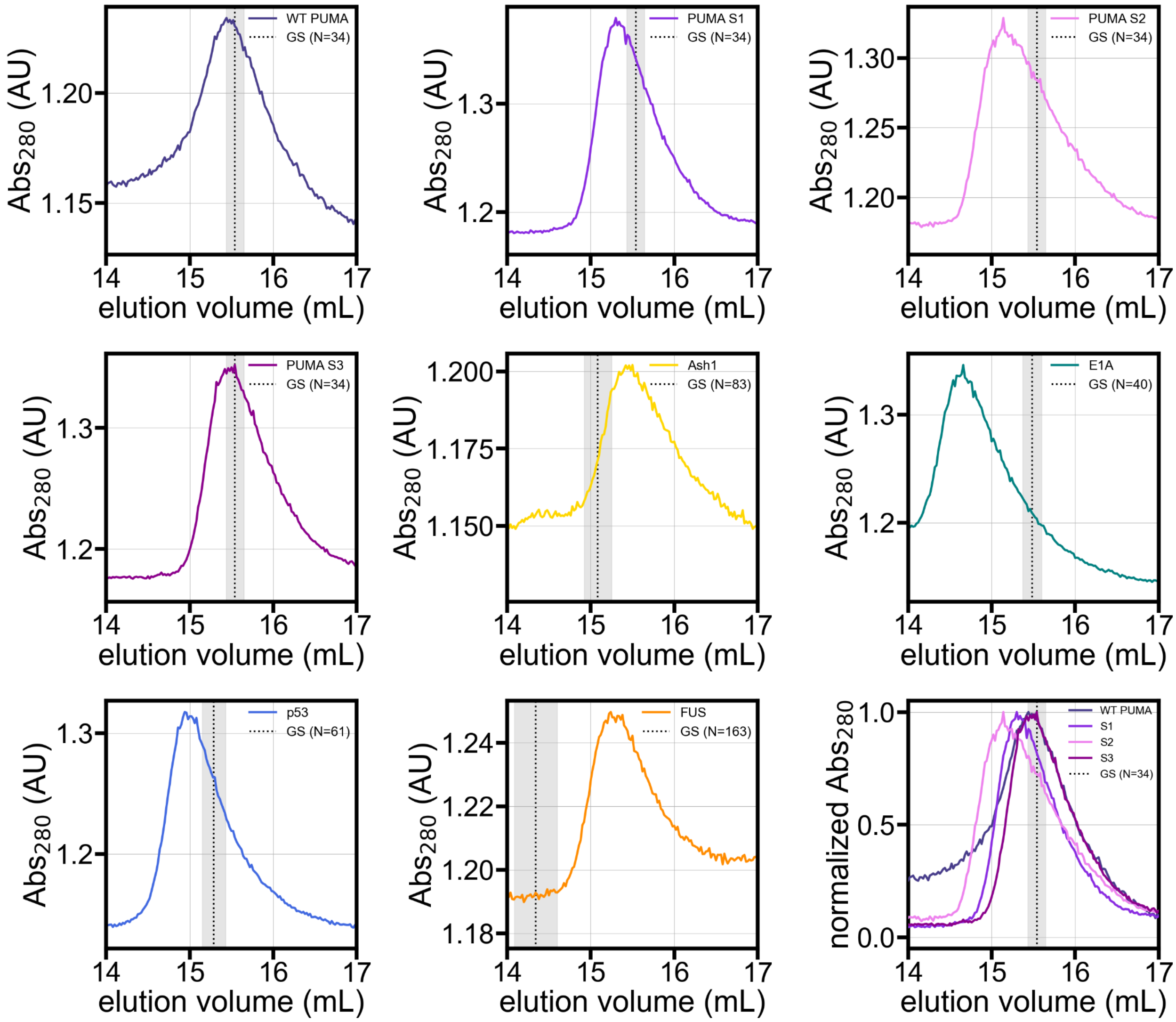


**Figure S2.** Chromatograms from SEC-SAXS experiments in which the samples were donor-IDP-acceptor FRET constructs in a dilute phosphate buffer solution. Vertical dotted line labeled “GS” in each panel represents the expected elution peak position of a FRET construct containing a GS-repeat sequence equal in length to the IDP, where N refers to the number of amino acids. Shaded region in each panel represents the standard error of the expected GS peak position.

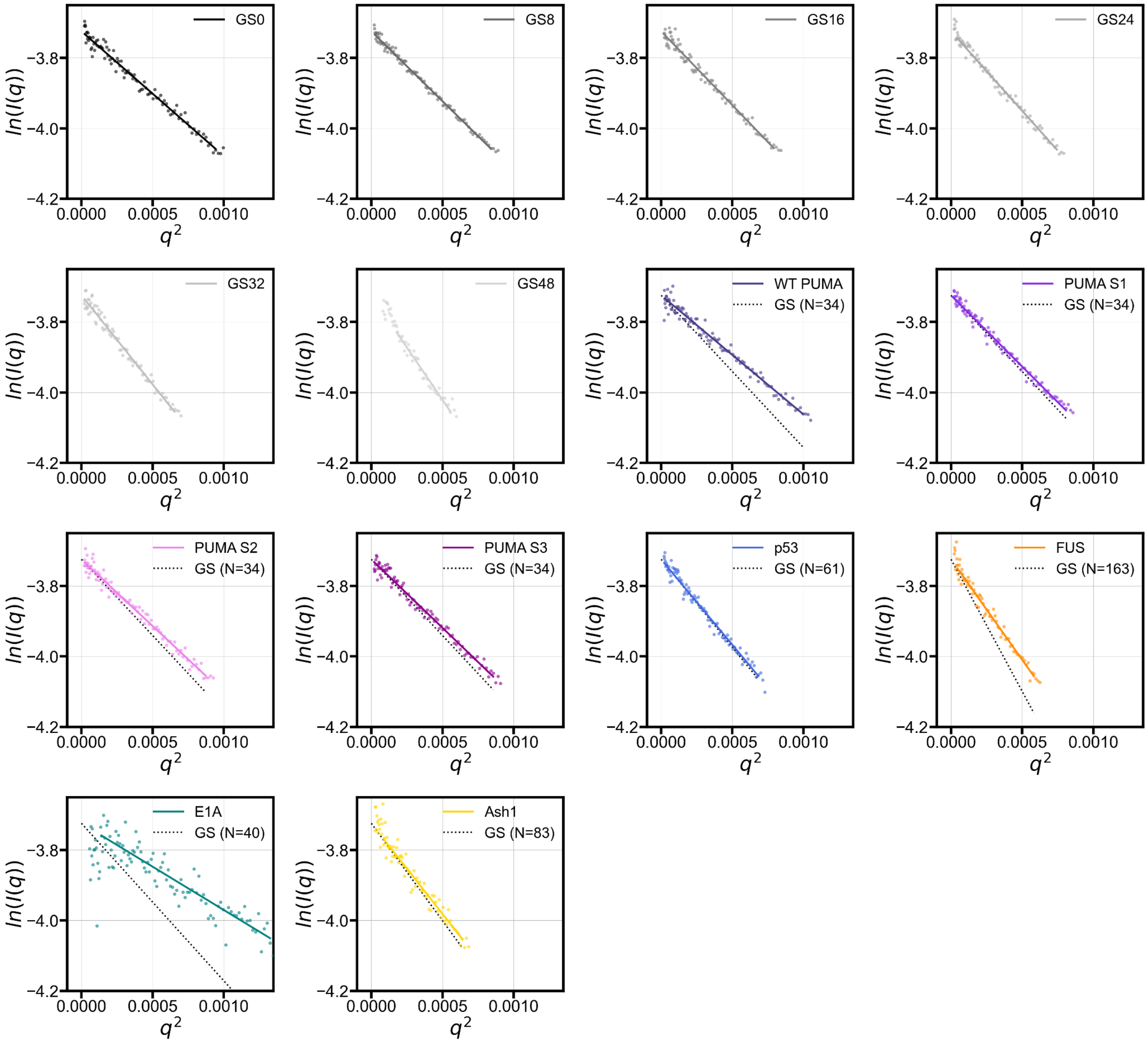


**Figure S3.** Guinier plots for donor-IDP-acceptor FRET constructs from SEC-SAXS experiments. For IDRs that are not GS-repeat sequences, the fitted line is compared with the expected fitted line for a construct containing a GS-repeat sequence of the same length (black dotted lines), where N refers to the number of amino acids. Lines are fitted to a maximum $q * R_g$ value of 1.

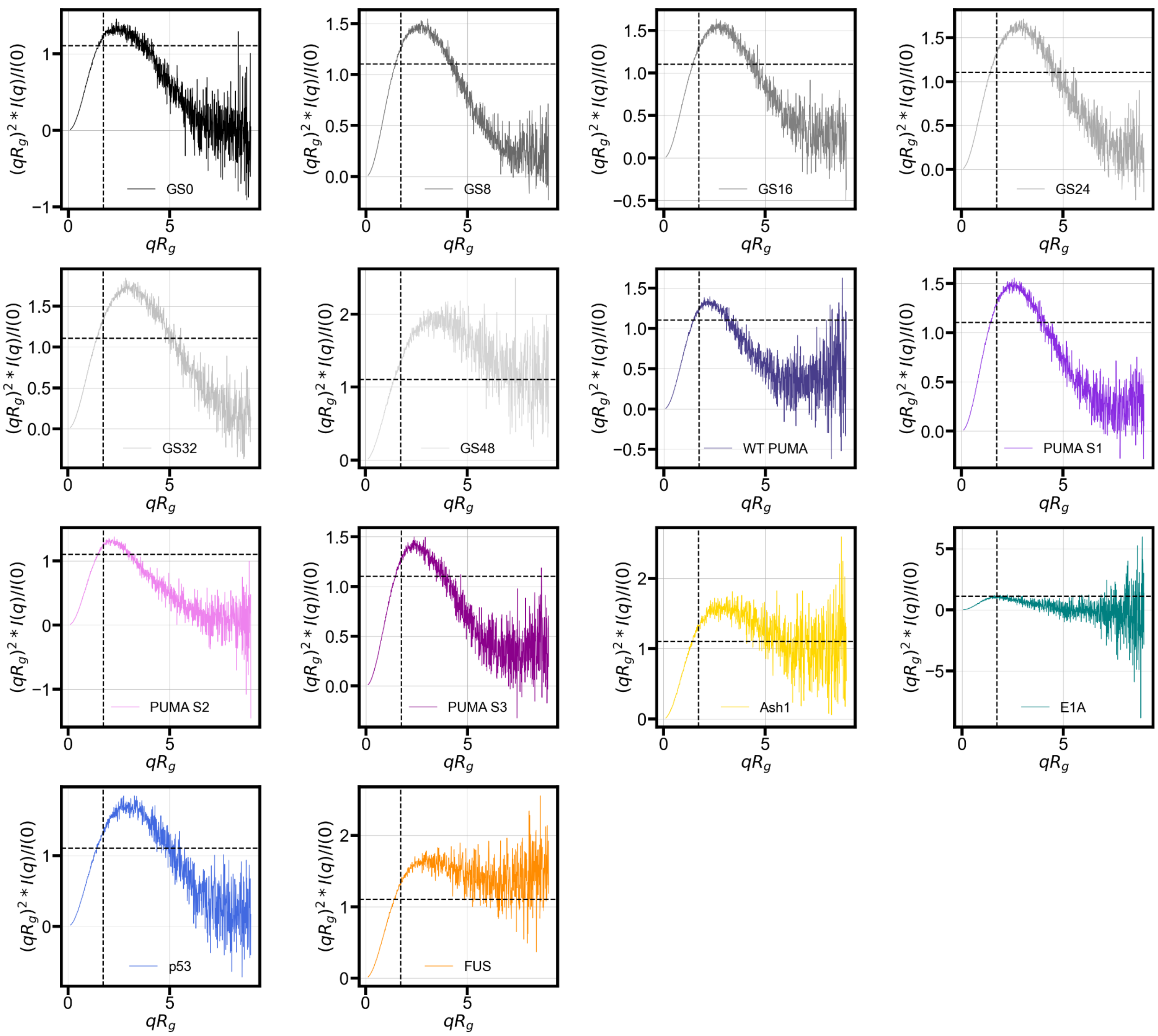


**Figure S4.** Dimensionless Kratky plots derived by transforming the scattering profiles from which the $R_g$ values reported in the main text were calculated. For a globular protein, the peak position should be at $qR_g = \sqrt{3} \sim 1.73$ (shown by vertical dashed line) and the peak height should be $(qR_g)^2 * I(q)/I(0) = 3/e \sim 1.1$ (shown by horizontal dashed line).

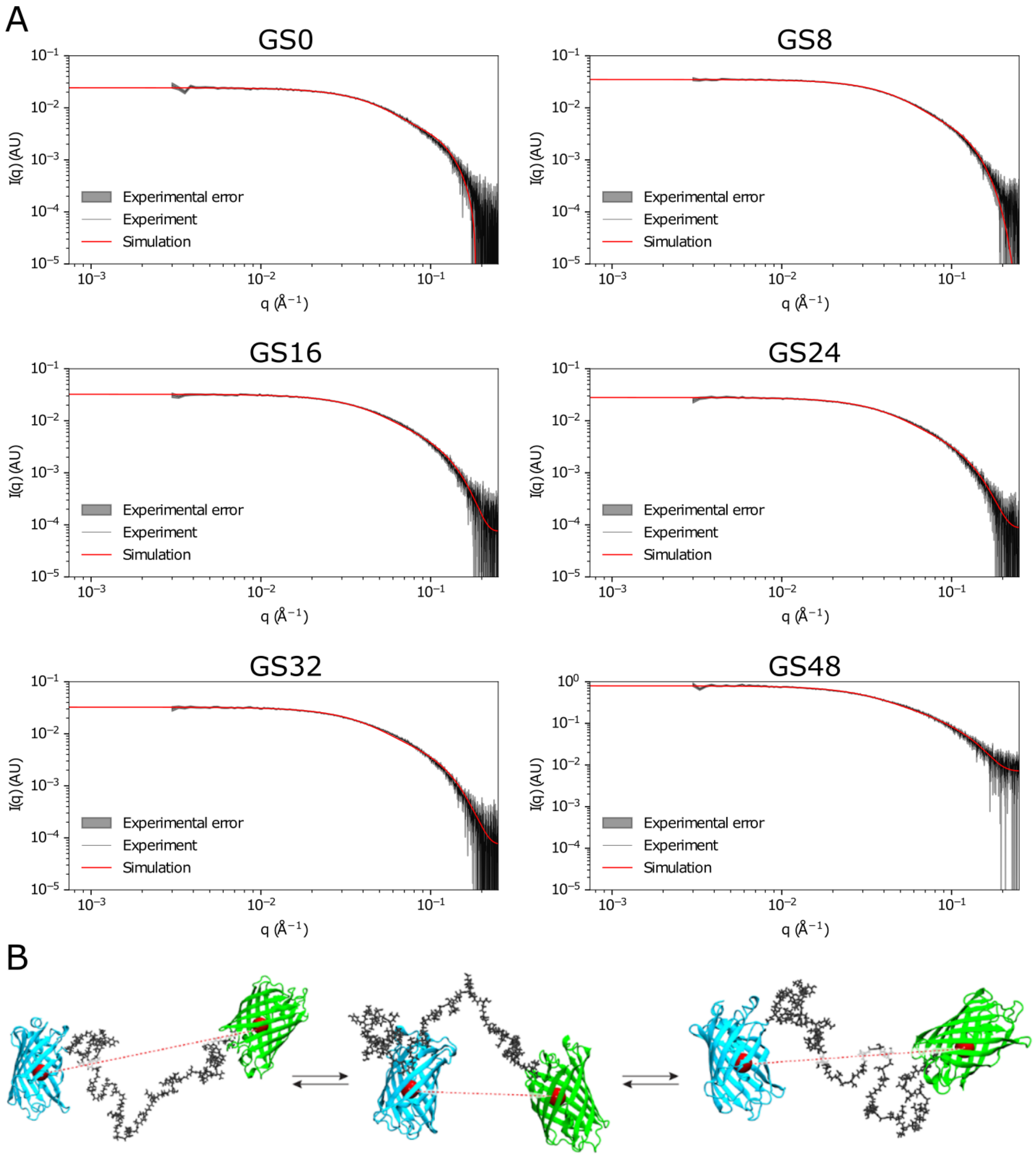


**Figure S5.** (**A**) Comparison of experimentally measured small-angle X-ray scattering profiles with simulation-derived synthetic for GS-repeat sequences of different lengths. (**B**) Example snapshots from simulations. $R_e^{app}$ is calculated based on the distance between residues from the center of the two beta-barrels (dashed red line).

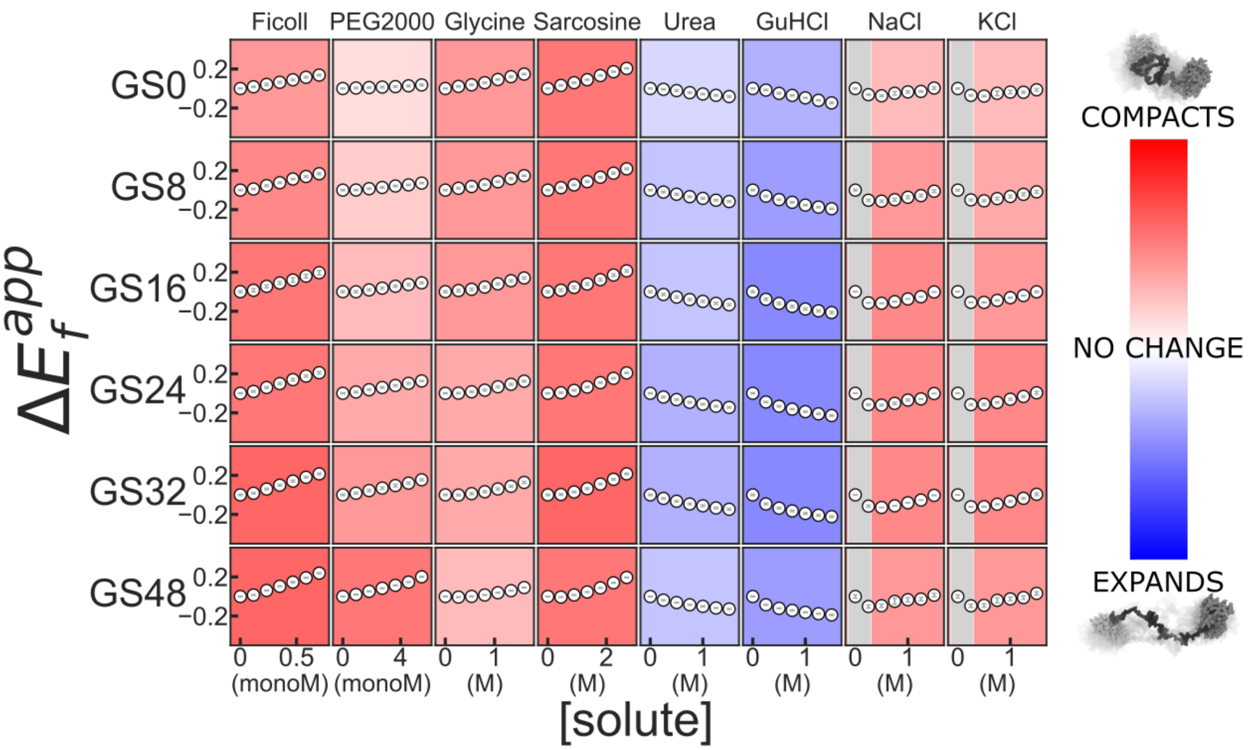


**Figure S6.** Solution space scans of GS-repeat homopolymers. Each cell shows $\Delta E_f^{app}$ as a function of increasing solute concentration. Blue background indicates expansion and red indicates compaction, with deeper colors indicating more change. monoM: Concentration of a polymer expressed as a concentration of monomeric units. Light gray shaded regions on left side of cells for solutes NaCl and KCl: approximate range of concentrations within which electrostatic screening is the dominant effect; the leftmost two points of each series, since they are within that range, are not used in the assignment of background color.

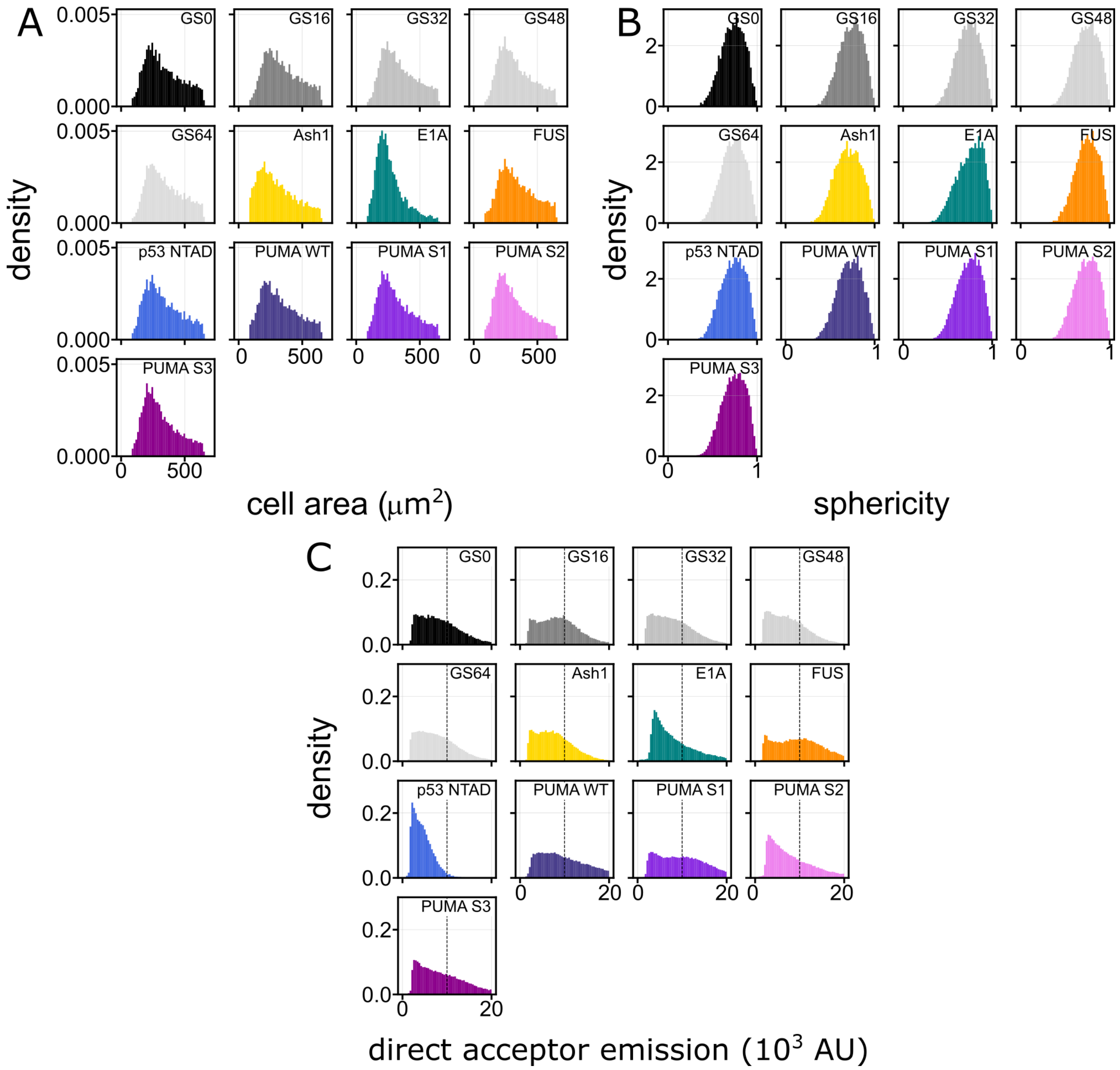


**Figure S7.** Probability density of cellular features across cells expressing different FRET constructs. Each histogram contains over $10^3$ cells, with a total of over $2.1 \times 10^5$ cells. (**A**) Cell area (in $\mu m^2$). (**B**) Cell circularity calculated as $4\pi(area/perimeter^{2})$. A value of 1 is a perfect circle and as the value approaches 0 it indicates an increasingly elongated polygon. The HEK293T cells used in these experiments tend to be rounder than others, resulting in a sphericity close to 1. (**C**) Direct acceptor emission (cells excited at 511 nm) is a metric for in-cell construct concentration. To eliminate artifacts resulting from high overexpression, and to facilitate accurate comparison with *in vitro* measurements done at 1 uM, we left cells with a direct acceptor emission higher than 10,000 out of the analysis, indicated by the dashed vertical line (see also **Fig. S17**).

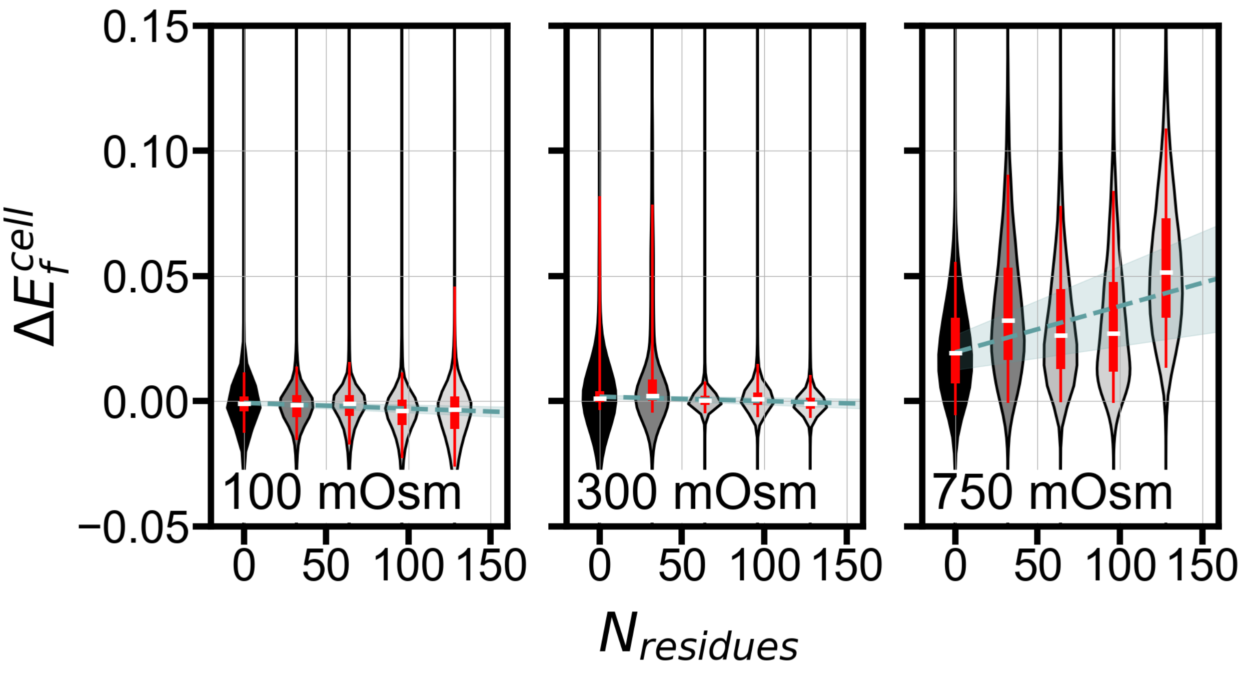


**Figure S8.** Linear fit of $\Delta E_f^{cell}$ as a function of GS linker length for hypo, iso, and hyperosmotic perturbations. Dashed green line is a linear fit of the medians, shown as white dashes, and shaded areas are the errors of the fit. Thick and thin red bars span median 50% and 90% of the data, respectively. For the number of cells used to generate each violin plot, see **Table S3**.

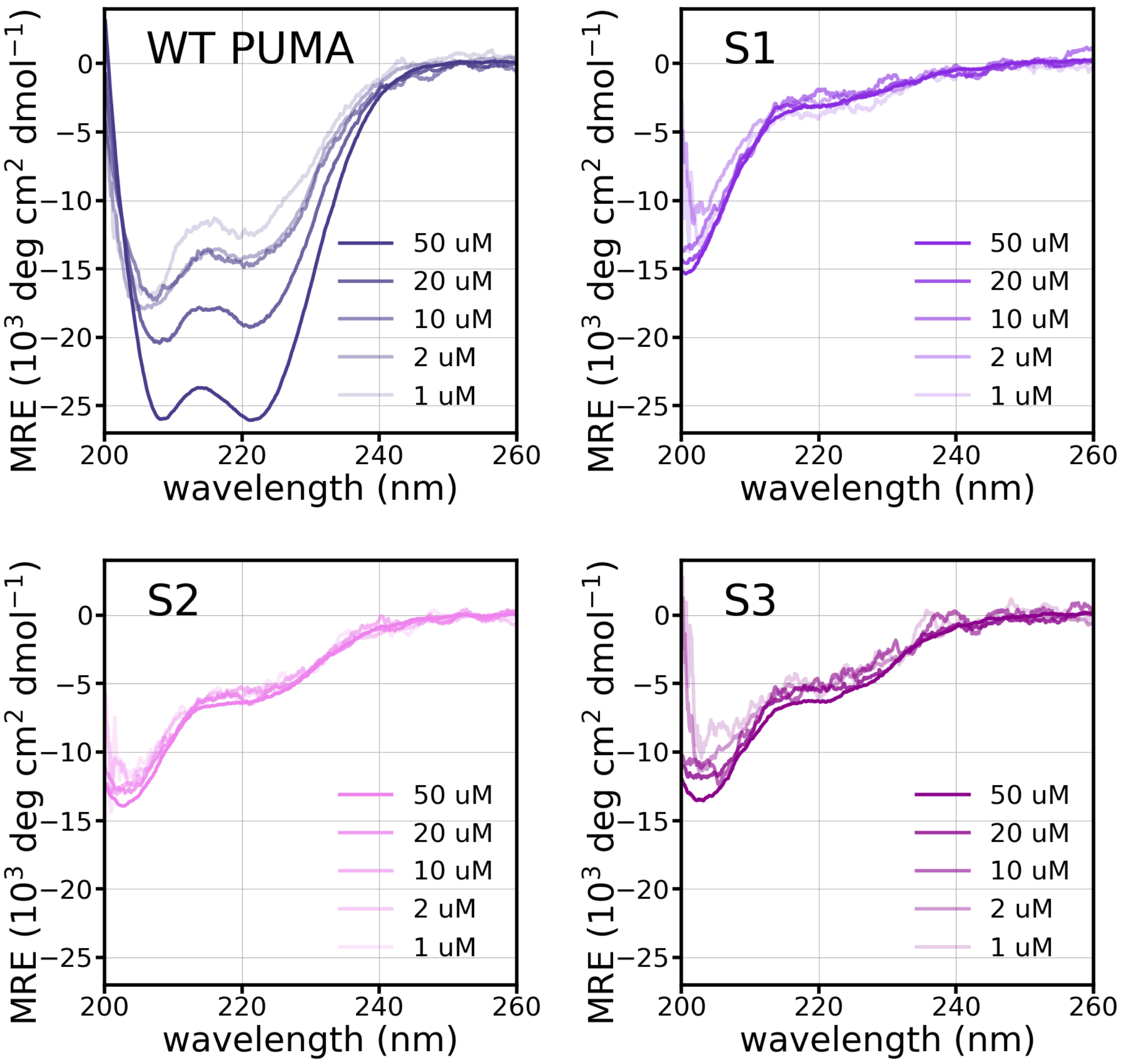


**Figure S9.** Concentration dependence of circular dichroism measurements of PUMA WT and sequence scrambles.

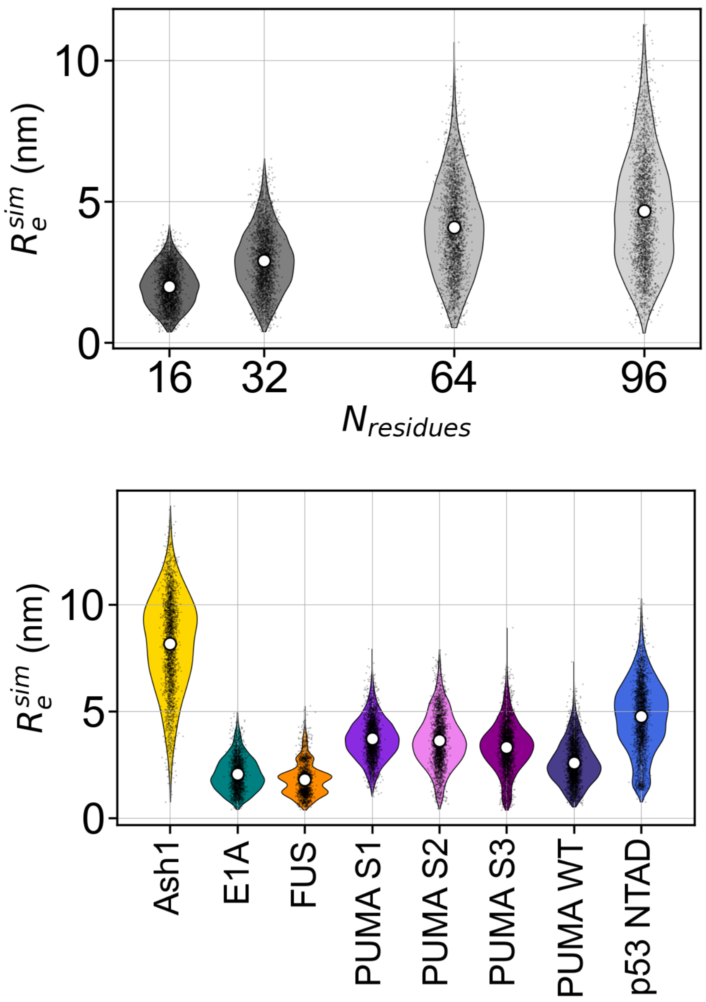


**Figure S10**. All-atom simulations, using the ABSINTH forcefield[2], of all sequences in this work. Simulations did not contain the fluorescent protein labels. Violin plots are obtained from random sampling of 2000 frames from ensembles containing at least 20,000 conformations. White circles represent the mean and black points are individual frames. FUS shows a multimodal distribution since its ensemble is poorly sampled, and may not be indicative of a well-sampled ensemble for this construct.

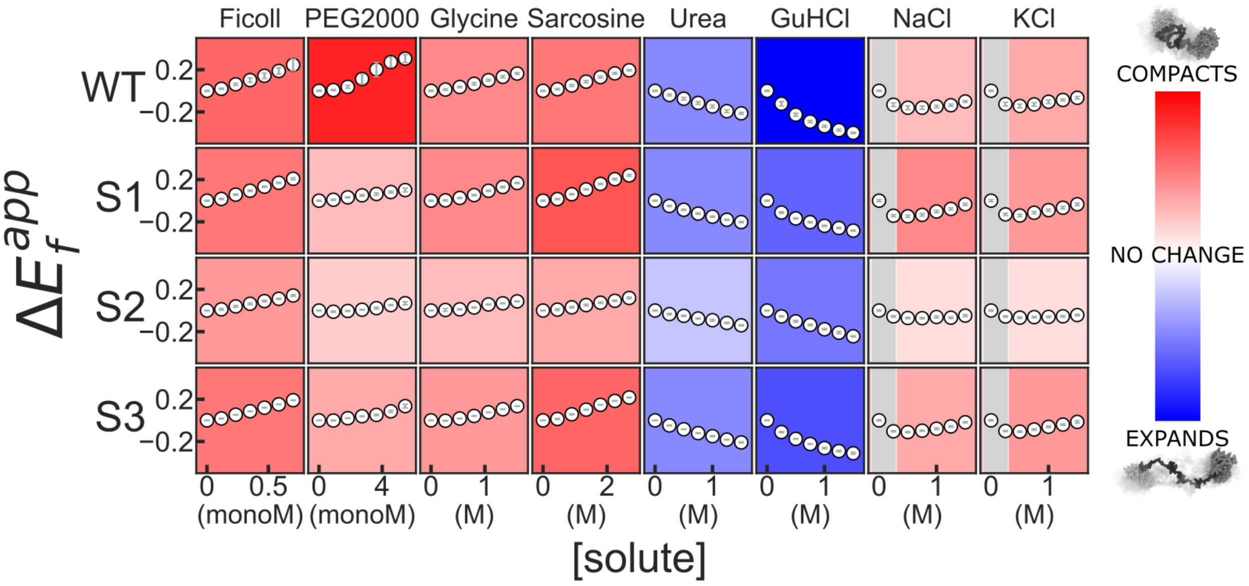


**Figure S11.** Solution space scans of WT PUMA and sequence scrambles. Features are as in **Fig. S6**.

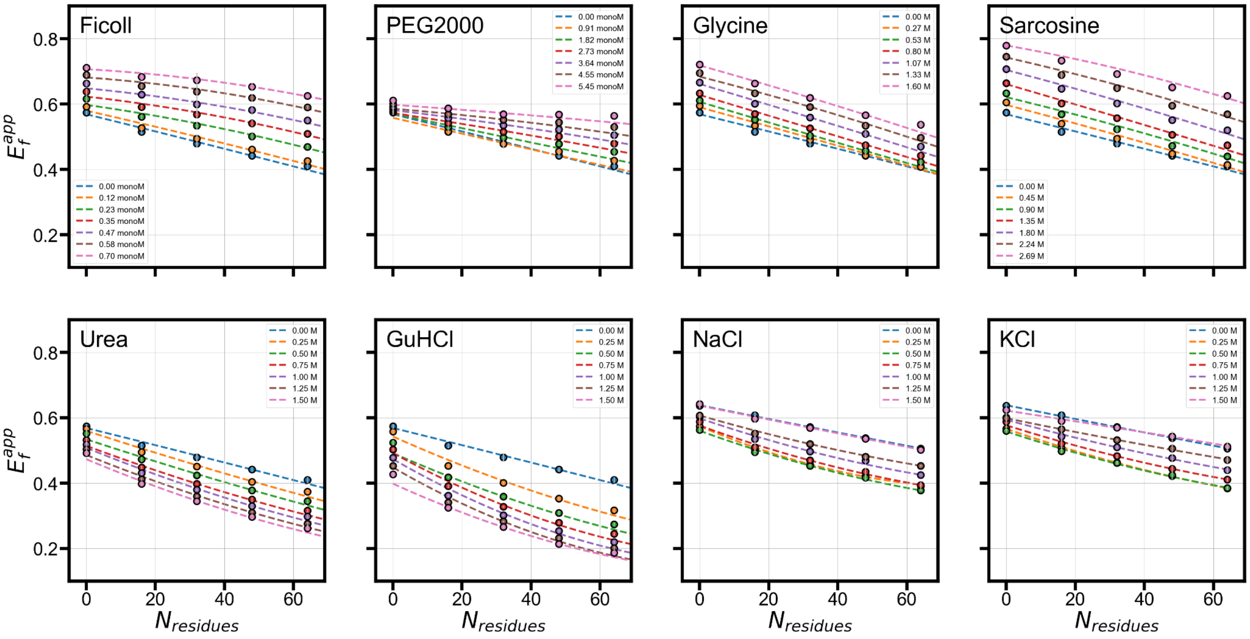


**Figure S12.** $E_f^{app}$ vs. length of GS-repeat sequence in various solution conditions. A second degree polynomial fit, shown as dashed lines, is used for interpolation of $\Delta E_f^{app}$ for arbitrary sequence lengths in **Figs. 3H** and **4E**.

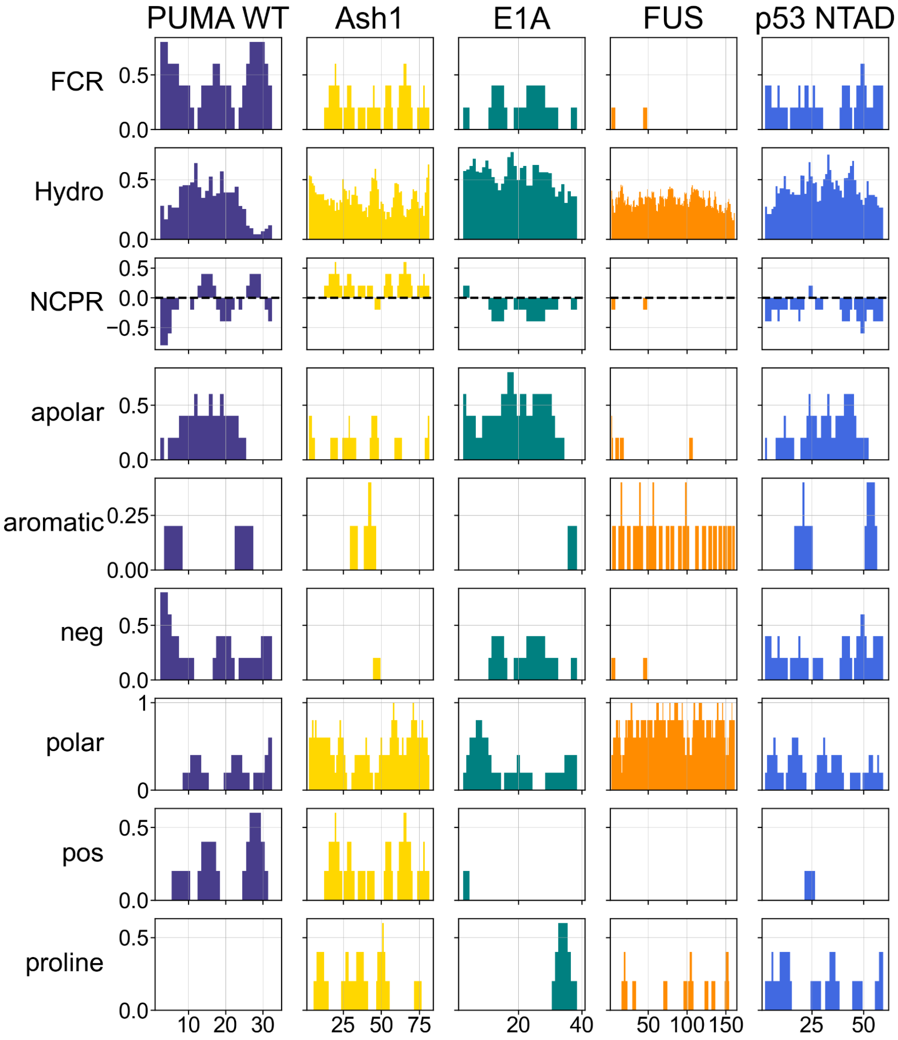


**Figure S13.** Sequence features of IDP sequences in **Fig. 4**, calculated using localcider[3]. All bars represent the average over a five-residue window centered at the specified residue number. FCR: Fraction of charged residues; Hydro: Kyte-Doolitle hydrophobicity scale; NCPR: net charge per residue; apolar: fraction of ALMIV residues; aromatic: fraction of FYW residues; neg: fraction of ED residues; pos: fraction of KR residues; polar: clusters of QNSTGHC residues; proline: fraction of P residues.

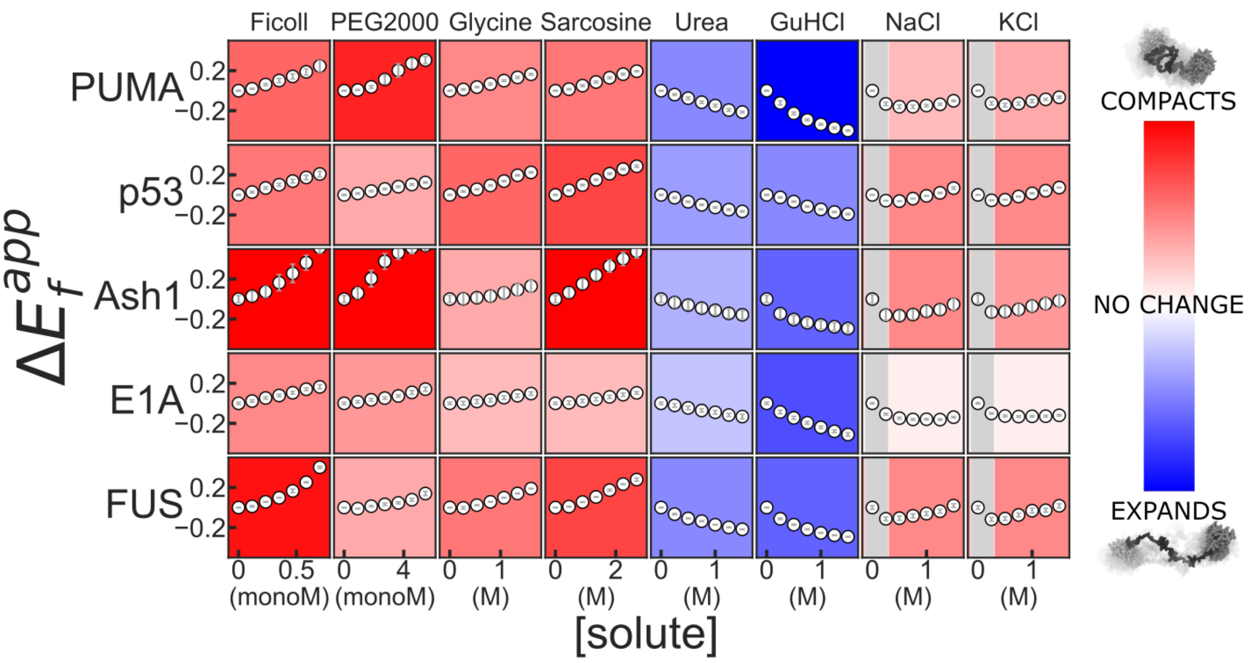


**Figure S14.** Solution space scans of naturally occurring IDRs. Features are as in **Fig. S6**.

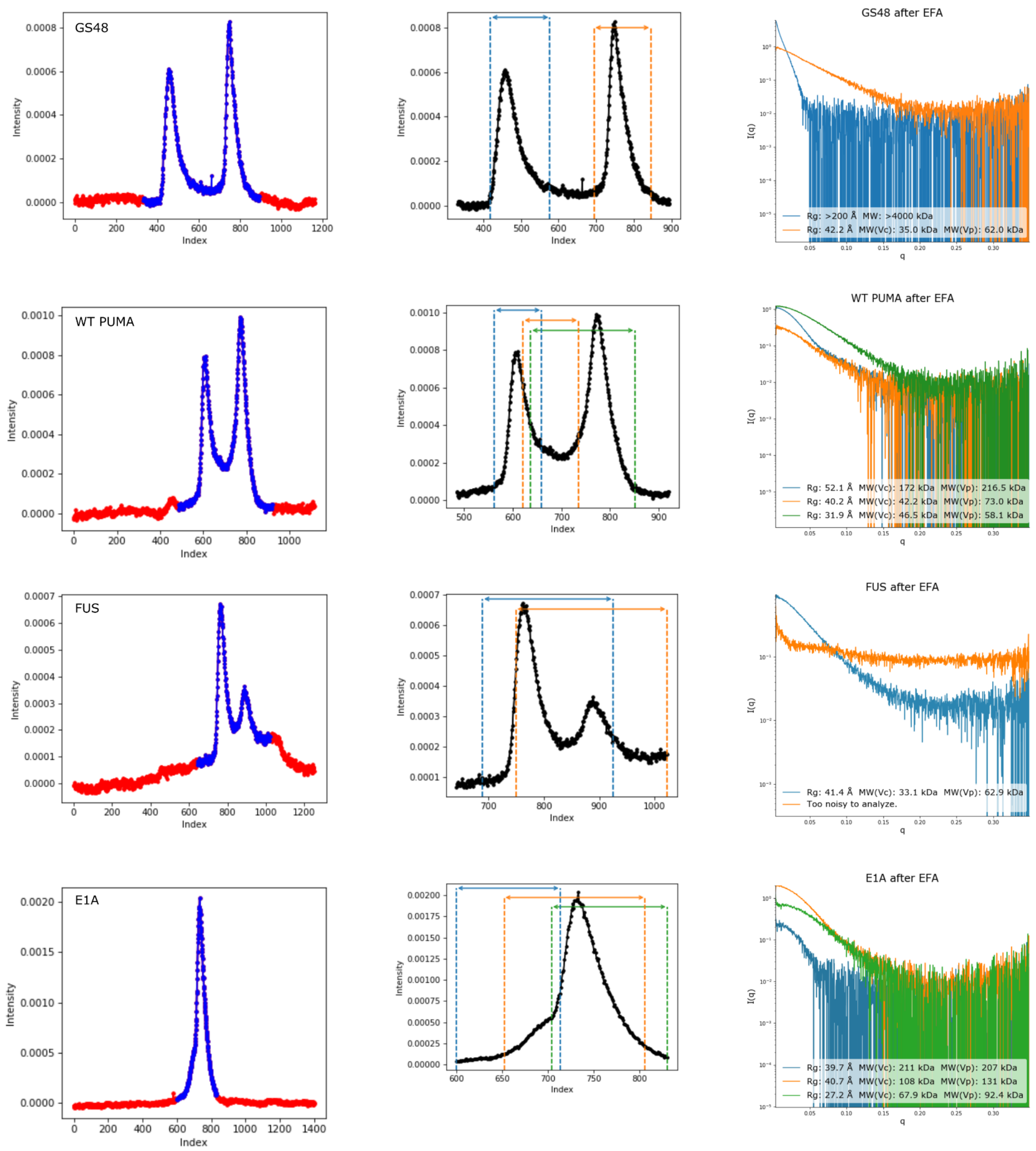


**Figure S15.** Screens from BioXTAS RAW software[1] showing process of deconvolution of SEC peaks using evolving factor analysis. Left: raw chromatograms. Center: ranges of deconvoluted peaks. Right: $I(q)$ *vs.* $q$ series, calculated radius of gyration, and calculated molecular weight for each deconvoluted peak. Same colors in center and right panels represent the same deconvoluted peaks.

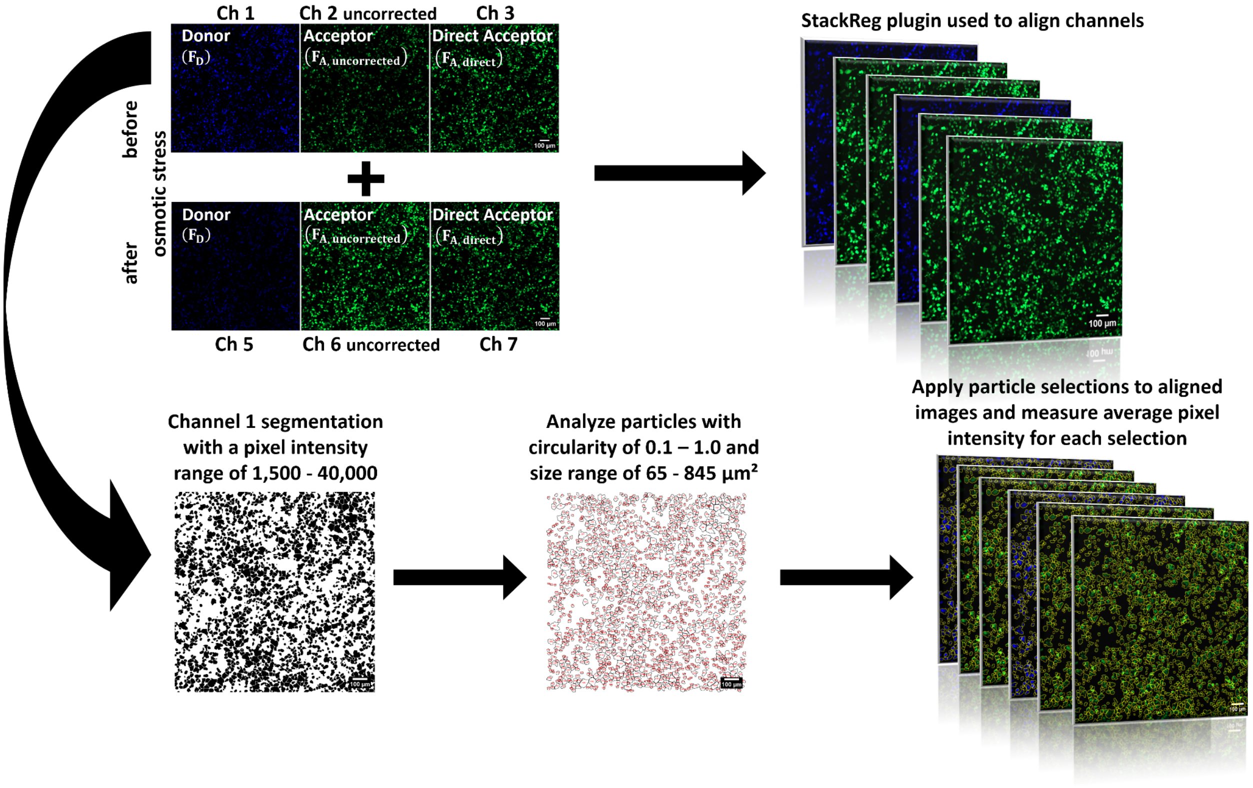


**Figure S16.** Analysis pipeline for live cell data. The donor channel before perturbation (Ch 1) was segmented using a fixed threshold to include any pixels with an intensity value between 1,500 - 40,000. The ImageJ "analyze particles" algorithm was used to select thresholded regions with a circularity between 0.1 -1.0 and a size of 65 - 845 µm². All channels were aligned using the StackReg plugin before segmented regions were applied and measured. Final measurements were corrected for bleedthrough and cross-excitation using slopes obtained from **Fig. S18.** The complete dataset can be found in **Table S2**. In this table, channels 1, 2, 2 uncorrected and 3 correspond to donor ($F_D$), corrected acceptor ($F_{A,}$), uncorrected acceptor ($F_{A, uncorrected}$) and direct acceptor ($F_{A, direct}$) before osmotic stress, respectively. Channels 5, 6, 6 uncorrected and 7 correspond to donor ($F_D$), corrected acceptor ($F_{A,}$), uncorrected acceptor ($F_{A, uncorrected}$) and direct acceptor ($F_{A, direct}$) after osmotic stress, respectively.

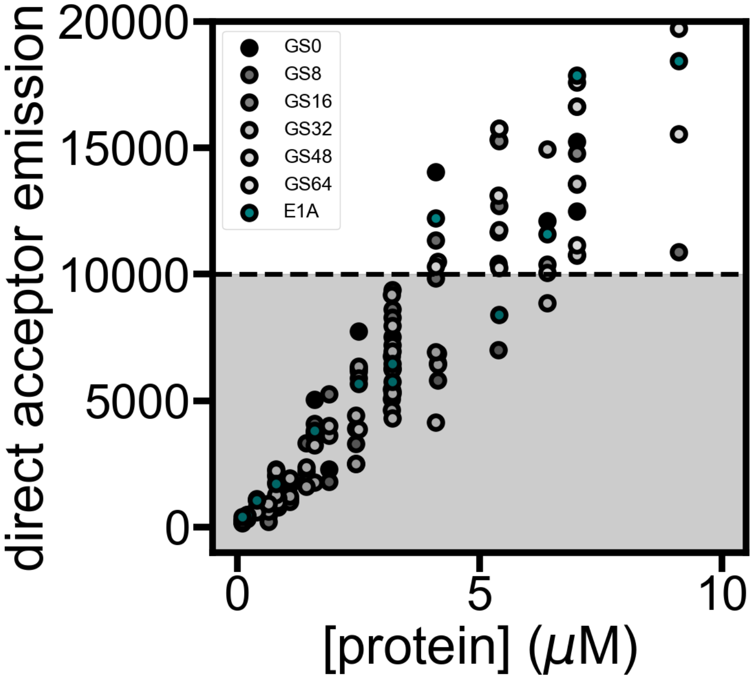


**Figure S17.** *In vitro* measurement of direct acceptor emission for known recombinant, purified proteins measured on the same setup as the live cells. Dashed line shows the emission cutoff used to select cells with a concentration range around 5 μM or lower to correlate with *in vitro* experiments.

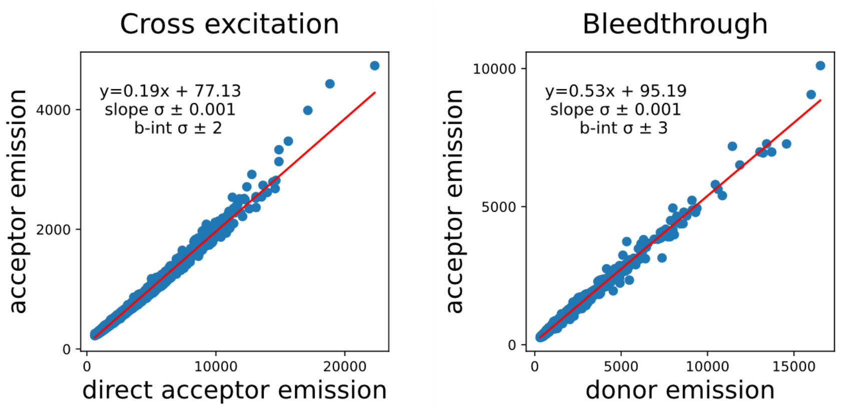


**Figure S18.** Measurements of cross-excitation (left) and bleedthrough (right) from donor to acceptor channel. To calculate cross-excitation, cells expressing mNeonGreen only were imaged. To calculate bleedthrough, cells expressing mTurquoise2 only were imaged. In both cases, the same imaging settings as those used for FRET constructs were used. (left) The x-axis shows acceptor emission under acceptor excitation. (right) The x-axis shows donor emission under donor excitation. In both figures, the y-axis shows acceptor emission under donor excitation. The slopes of these two values were used to correct the signal from the FRET construct according to the following equation:

$$F_A = F_{A,uncorrected} - (0.19 \times F_{A,direct} + 0.53 \times F_D)$$

where $F_A$ is used to calculate $E_f^{cell}$. The numbers 0.19 ± 0.001 and 0.53 ± 0.001 are the slopes from the figures above.

Additionally, we performed photobleaching experiments where mNeonGreen of various FRET constructs were bleached. These bleached constructs were used to measure and calculate bleedthrough and similar results were obtained (slope of 0.51 ± 0.007).

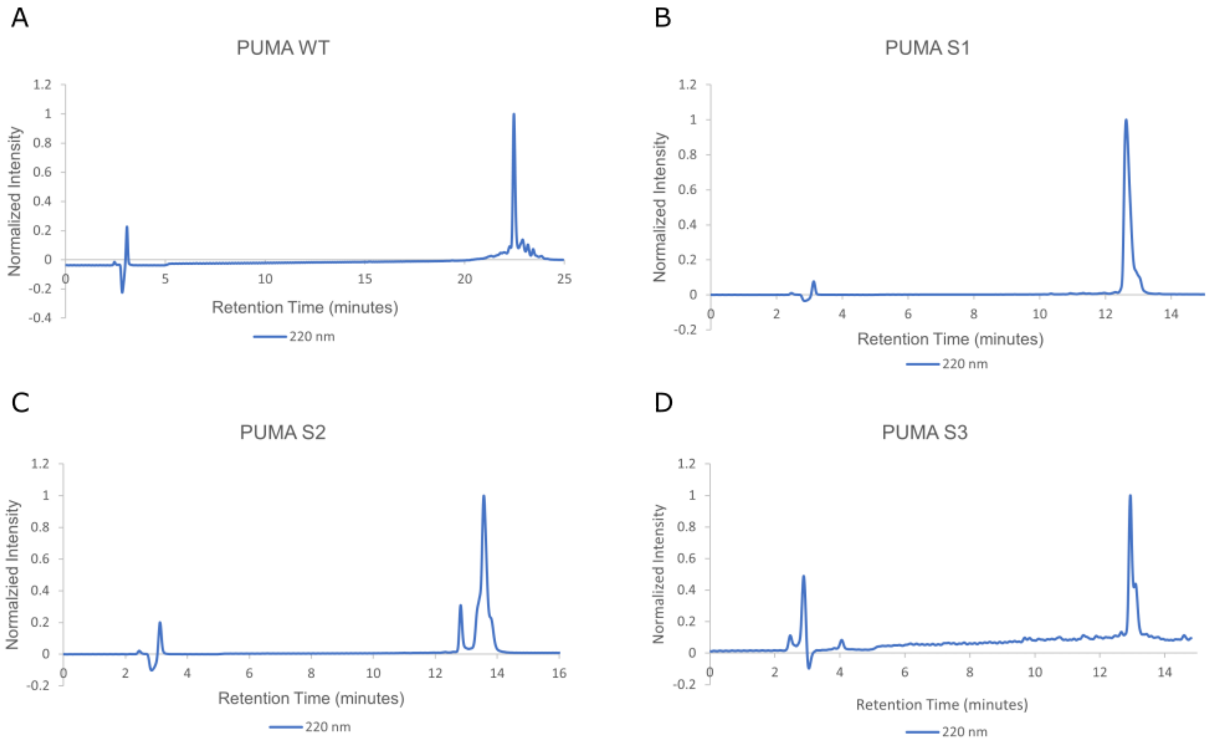


**Figure S19.** HPLC traces from purification of label-free peptides. **(A)** PUMA WT. **(B)** PUMA S1. **(C)** PUMA S2. **(D)** PUMA S3.

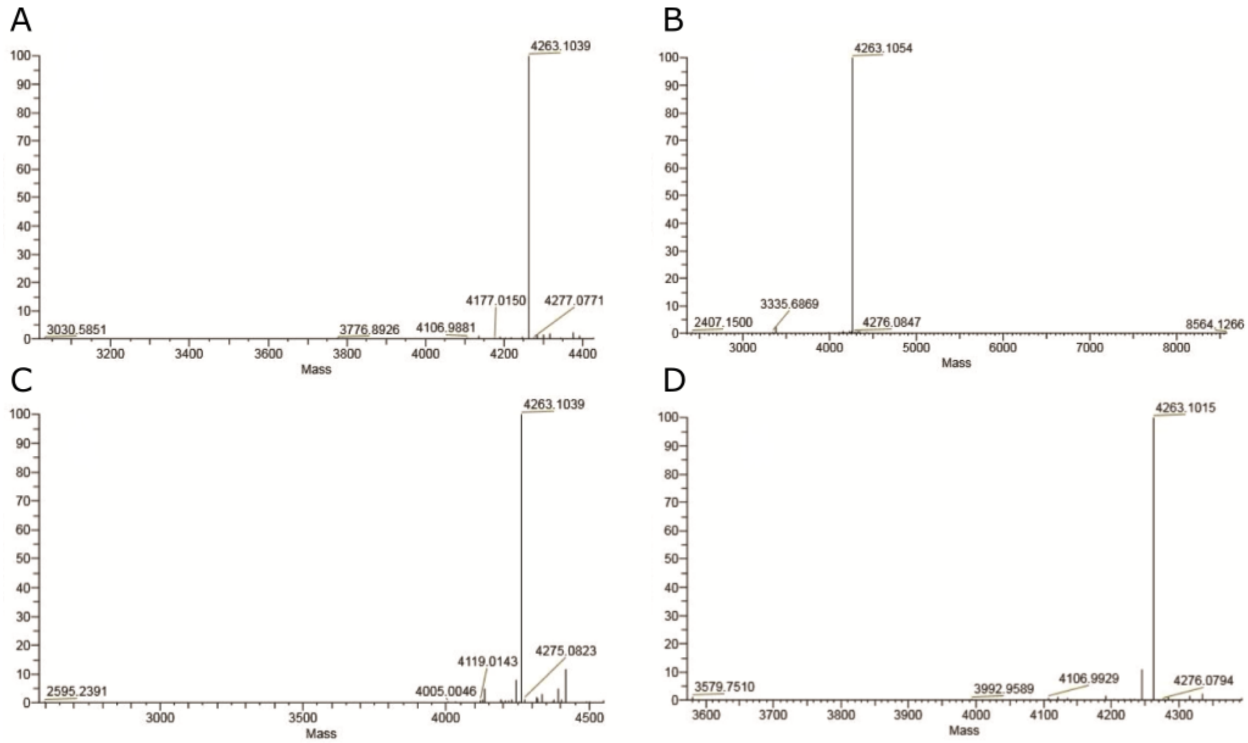


**Figure S20.** High-resolution ESI mass spectra of purified label-free peptides. (A) PUMA WT. (B) PUMA S1. (C) PUMA S2. (D) PUMA S3. Calculated and experimental masses are shown in **Table S4**.

| Peptide | Calculated Mass (Da) | Experimental Mass from ESI-MS (Da) | Retention Time (minutes) |
|---|---|---|---|
| PUMA WT | 4263.0918 | 4263.0974 | 22.47 |
| PUMA S1 | 4263.0918 | 4263.1014 | 12.64 |
| PUMA S2 | 4263.0918 | 4263.0923 | 13.59 |
| PUMA S3 | 4263.0918 | 4263.0948 | 12.94 |

**Table S4.** Calculated and experimental masses of label-free peptides.
